# A kinome inhibitor screen implicates adhesion and growth factor signaling in cellular recovery after caspase activation

**DOI:** 10.64898/2026.01.06.697929

**Authors:** Maddalena Nano, Jacob Harwood, Gabriel Lukaszewicz, Alexander Kagan, Leonid Peshkin, Denise J. Montell

**Affiliations:** Molecular, Cellular, and Developmental Biology Department, University of California, Santa Barbara, CA 93106; Neuroscience Research Institute, University of California, Santa Barbara, CA 93106; Department of Statistics, University of Michigan, Ann Arbor, MI 48109; Harvard Medical School, Systems Biology, 200 Longwood Ave, Boston MA 02115

## Abstract

Apoptosis is a common form of regulated cell death and requires cysteine-aspartic proteases called effector caspases. Caspase activation triggers positive feedback, leading to the idea that apoptosis is irreversible. However, we and others have demonstrated that cancer cells can survive effector caspase activation and become more aggressive and drug-resistant as a result. Despite the profound implications of apoptotic reversal, also known as anastasis, for both regenerative medicine and cancer therapy, the molecular pathways that enable cells to survive executioner caspase activation remain largely unmapped. To systematically dissect this phenomenon, we developed a quantitative screening platform that combines inducible caspase activation with kinome-wide pharmacological profiling. This approach uniquely allowed us to: (1) identify pharmacological modulators of post-caspase survival, (2) identify specific kinases regulating post-caspase survival, and (3) distinguish general toxicity from anastasis effects. This approach implicated regulators of cell adhesion and the cytoskeleton, consistent with the known rounding of apoptotic cells and respreading during recovery. Growth factor signaling also emerged from the analysis. In addition to its expected effects on unstressed cells, fetal bovine serum markedly increased anastasis. Some growth factor combinations were more effective than individual ones at recapitulating the serum effect. Similarly, pleiotropic kinase inhibitors were generally more effective than selective ones. Nevertheless, selective Rho kinase inhibition significantly enhanced anastasis whereas Akt inhibition impaired it, suggesting that these kinases serve as central nodes that integrate multiple upstream inputs. Beyond identifying specific kinase targets, our work provides a framework for ‘anti-anastasis’ therapies that could prevent cancer cell recovery after chemotherapy.

## Introduction

Intra- or extracellular stressors, as well as ligands such a tumor necrosis factor alpha, can trigger apoptotic pathways; yet cells do not invariably die (1–10). For example, cardiomyocytes (11) and neurons (12) can remain viable even after activation of effector caspases, which are the proteases that normally execute apoptosis (1,5,13). Thus, cellular survival after potentially lethal stress, also known as anastasis, can preserve terminally differentiated cells. Such forms of cellular resilience also promote tissue repair and regeneration. In irradiated *Drosophila* eye discs, where the vast majority of cells activate apoptotic caspases, those that survive contribute to regeneration (14). Moreover, sublethal caspase activation also enhances skin and corneal wound healing in mice, supports tail regeneration in tadpoles, and promotes axon regrowth in *C. elegans* (15). However, survival of damaged cells can also have detrimental consequences. Sublethal treatments can induce genomic instability (16), promote carcinogenesis in the skin (17), and generate drug-tolerant persister cells (18). Survival after damage can promote a more aggressive phenotype in melanoma cancer cells (19), drive ovarian cancer progression (20), and increase metastasis and chemoresistance in colorectal cancer cells (21), as well as exacerbate the malignant behavior of breast cancer cells (22). The apoptotic machinery is also co-opted by cells that survive oncogene-targeted therapies to resume proliferation (23). Therefore, it is critical to identify mechanisms that determine whether cells live or die in response to damage.

To identify pharmacological modulators and gain insight into the mechanisms enabling cells to recover from caspase activation or inhibiting their survival, we developed a cell line that allows us to induce and simultaneously monitor executioner caspase activity (24). We engineered HeLa cells to encode a doxycycline (DOX)-inducible caspase-3 (24,25) and a sensor for executioner caspase activity (GC3AI) (26). In the presence of DOX, active caspase-3 is expressed, the sensor is cleaved, and cells fluoresce. Green fluorescence persists until the sensor turns over or is diluted by cell division.

Protein kinases regulate most cellular processes including apoptosis and anastasis (20,27–36). Moreover, kinases are highly druggable targets, with 80 FDA-approved inhibitors in clinical use (37). Many kinase inhibitors are pleiotropic, which can be exploited because each drug has a unique constellation of targets, and each kinase exhibits a specific pattern of inhibition that can serve as a molecular fingerprint. Thus, a relatively small number of kinase inhibitors can interrogate most of the 518 human protein kinases. Prior knowledge of how each inhibitor affects each target (38,39) is then used to implicate specific kinases in a process named kinome regularization (39,40) (**Figure 1A-B**). For example, AKT2, CDK6, CAMKL2, and PKACG were identified as regulators of endothelial cell proliferation in response to FGF2, VEGFA, and HGF (39,40), and MINK1, the SRC family kinases Fyn, Lyn, and YES1, and the ephrin receptor EPHB2 contribute to cell migration in a wound-healing assay (39,40). Alternatively, if there is substantial functional redundancy amongst individual kinases, promiscuous inhibitors may be identified that enhance or inhibit a process that more specific inhibitors might fail to affect.

**Figure 1.**
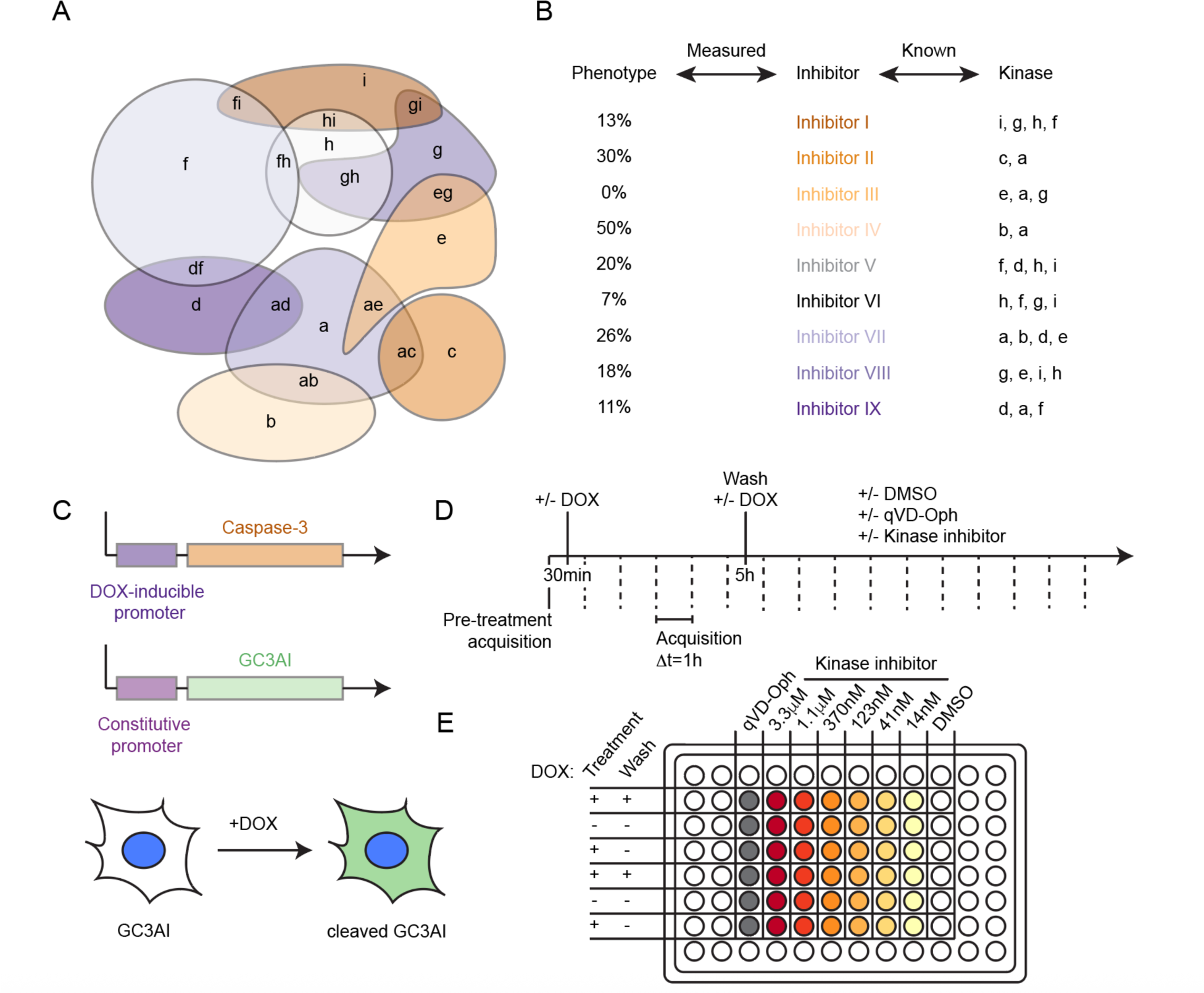
Identifying pleiotropic kinase inhibitors that modulate cellular resilience. **A.** A Venn diagram illustrating the concept of overlapping pleiotropies of kinase inhibitors (represented by different colors) and their targets (a-i). **B.** Each kinase has a fingerprint of inhibition. **C.** Genetic makeup of HeLa iCasp3-GC3AI cells. **D-E.** Screen timeline (D) and plate layout (E). Cells were treated +/- DOX for 5h, and were then washed in media +/- DOX containing DMSO or the caspase inhibitor qVD-Oph, or one of 6 concentrations of a kinase inhibitor (n=2).

Using this method, we identified 4 pleiotropic kinase inhibitors that enhanced survival after transient caspase activation at multiple concentrations, and 12 that potently inhibited it. Our findings reveal three fundamental principles of cellular resilience: first, anastasis depends on redundant signaling networks rather than single pathways. Second, classical pro-survival signals like growth factors and Akt remain critical even after caspase activation, extending their protective roles. Third, cytoskeletal regulators like ROCK function as molecular switches whose modulation can tip cells toward recovery or death. These insights reframe our understanding of cell fate decisions and identify new targets for therapeutic intervention.

## Results

### 1. A screen for compounds that modulate cellular resilience

To control and monitor caspase activation, we used a HeLa cell line (HeLa iCasp3-GC3AI) that expresses a DOX-inducible form of activated caspase-3 and the GC3AI fluorescent reporter of caspase activity (see **Methods Section 1**) (24–26) (**Figure 1C**). We previously identified conditions that result in robust GC3AI activation (see methods) yet allow 70 to 85% of cells to survive (24).

To identify kinase inhibitors that modify the fraction of cells that survive caspase activation, iCasp3-GC3AI cells were mock-treated or exposed to an initial pulse of caspase activity (5h DOX) followed by treatment with one of 47 pleiotropic inhibitors previously determined to be most informative for the kinase regularization technique (39) (**Supplementary Table 1**), each at six different concentrations (3.3µM, 1.1µM, 370nM, 123nM, 41nM, 14nM). As a positive control, we treated cells with the irreversible caspase inhibitor qVD-Oph at a concentration that inhibits cell death but allows GC3AI activation. Inhibitors were either diluted in media to allow for recovery, or in media with DOX to induce cell death (**Figure 1D-E**).

After imaging cells hourly for ∼30h, we quantified anastasis by measuring the percentage of GC3AI^+^ cells that were alive in samples transiently exposed to DOX, and normalizing to the DMSO-treated control by subtraction (see **Methods Section 4.4**). Comparing the results across treatments, we identified 12 compounds that inhibited anastasis at all concentrations tested (Cdk1/2 inhibitor III, Vargatef, NVP-BGT226, Pluripotin, PF-3758309, AT9283, BMS-754807, PHA-848125, Dinaciclib, MGCD-265, Ponatinib, Lestaurtinib; **Figure 2**), 4 compounds that enhanced anastasis at multiple concentrations (JNJ-7706621, AT7867, PKI-SU11274, GSK-269962A; **Figure 3A-D**, **Table 1**), and 2 compounds that were neutral at all concentrations tested (GSK650394, Erlotinib hydrochloride, **Figure 3E-F**). Most of the remaining compounds exhibited dose-dependent toxicity in the presence or absence of caspase activity (**Supplementary Figure 1** and **2**).

**Figure 2.**
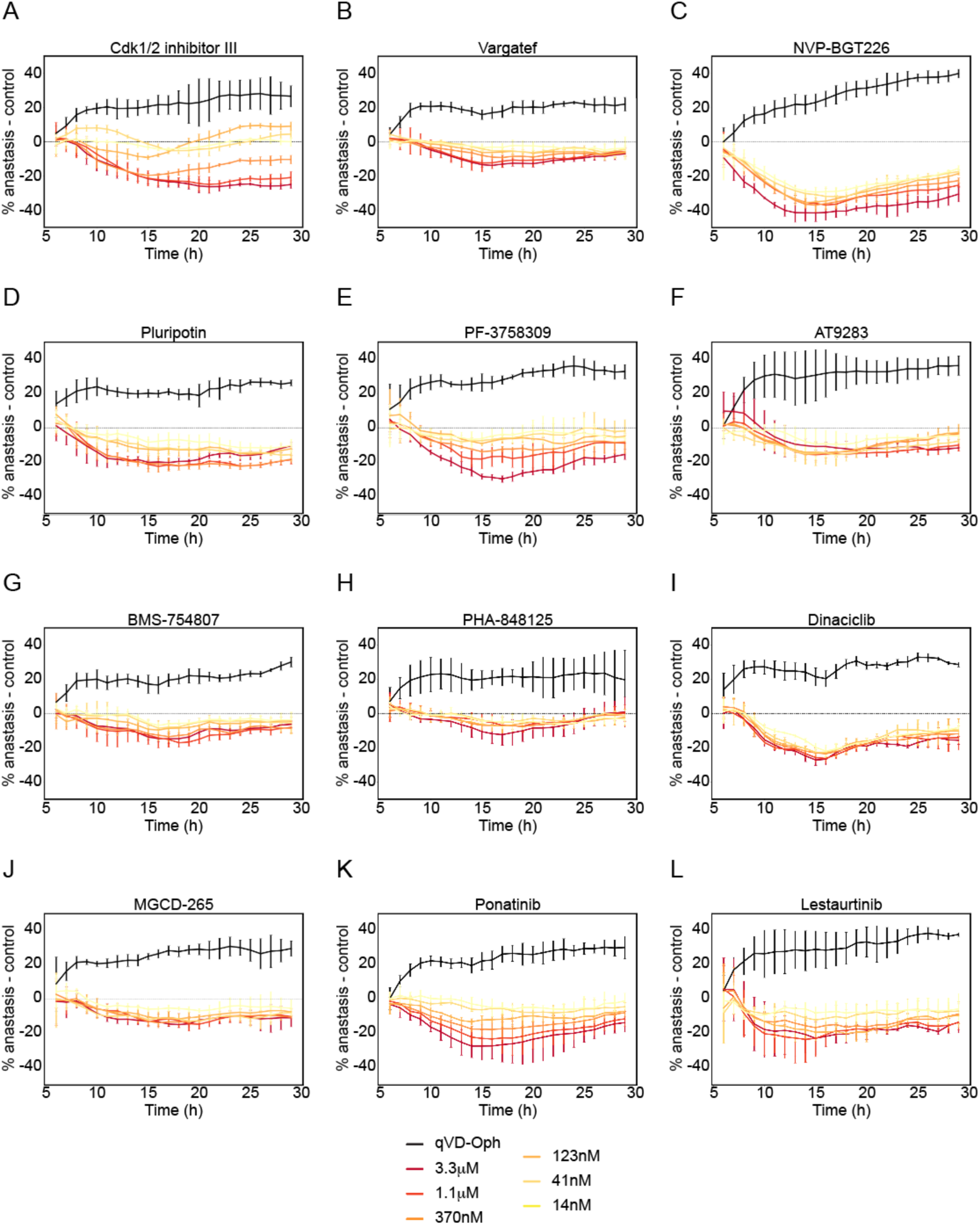
Compounds inhibiting anastasis at all concentrations tested. Percentage of cells undergoing anastasis relative to the control (DMSO) for compounds inhibiting survival following transient caspase activation (12–18h; see **Methods Section 4.5**). Data were normalized by subtraction, smoothed, and corrected as described in **Methods Section 4.8** and **4.9**. **A.** Cdk1/2 inhibitor III. **B.** Vargatef. **C.** NVP-BGT226. **D.** Pluripotin **E.** PF-3758309. **F.** AT9283 **G.** BMS-754807 **H.** PHA-848125. **I.** Dinaciclib. **J.** MGCD-265. **K.** Ponatinib. **L.** Lestaurtinib. Error bars represent the mean ± SD from n=2 independent experiments.

**Figure 3.**
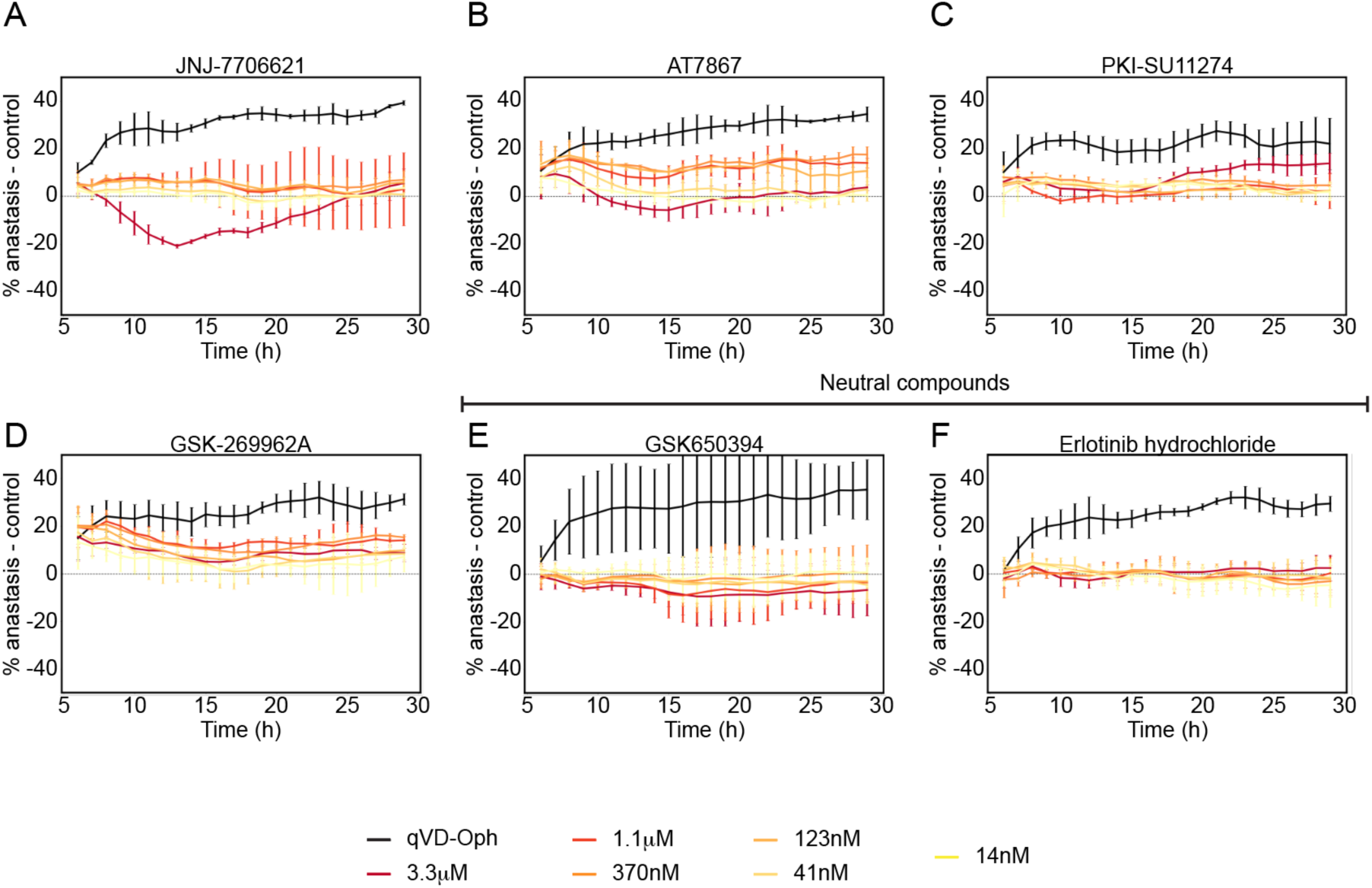
Compounds promoting anastasis. Plots showing the percentage of anastasis normalized to the control for compounds promoting survival following transient caspase activation at multiple concentrations (A-D) and for neutral compounds (E-F) (12–18 h; see **Methods Section 4.5**). Data were normalized by subtraction, smoothed, and corrected as described in **Methods Section 4.8**. **A.** JNJ-7706621. **B.** AT7867 **C.** PKI-SU11274 **D.** GSK-269962A. **E.** GSK650394. **F.** Erlotinib hydrochloride. Error bars represent the mean ± SD from n=2 independent experiments.

**Table 1.**
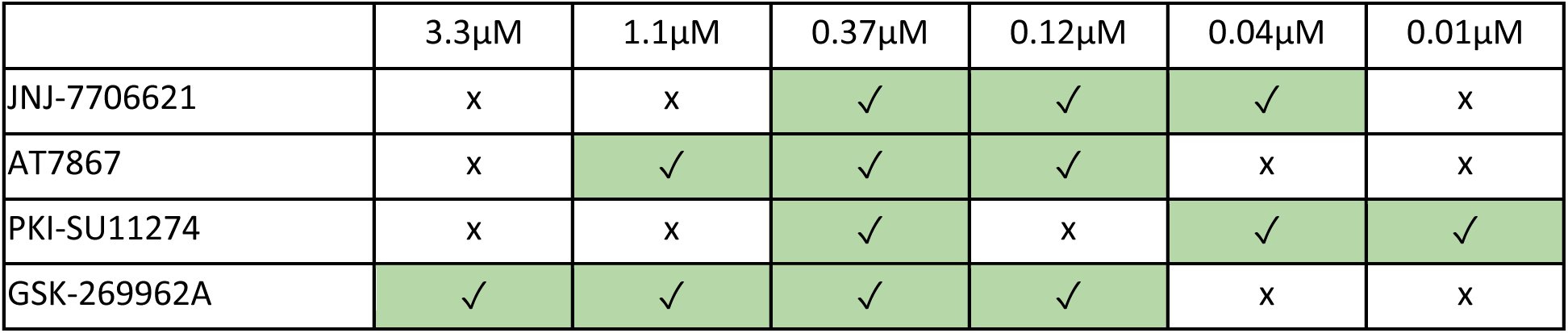
Kinase inhibitors that enhance survival after direct, transient caspase activation at multiple concentrations.

Several inhibitors, including those that enhanced anastasis, increased survival in DOX-treated, unwashed samples (**Supplementary Figure 1C**). This raised the possibility that some treatments might interfere with DOX-inducible gene expression rather than, or in addition to, affecting caspase-dependent recovery. To examine this, we generated a DOX-inducible mCherry cell line (**Methods Section 3**) and quantified how each of these inhibitors altered reporter expression (**Supplementary Figure 3**). We used these measurements to correct for effects on DOX induction (**Supplementary Figure 1**).

**Table 1**. Compound concentrations increasing anastasis (✓) (determined as described in **Methods Section 4.5**) are shown in green. All other concentrations are labeled with x.

### 2. Characterization of compounds inhibiting survival after direct caspase activation

We reasoned that compounds reducing survival might be equally toxic to unstressed and stressed cells, or they might more specifically inhibit anastasis. To distinguish between these possibilities, we compared cell survival in cells treated with each compound with or without DOX (**Methods Section 4.5** and **4.6**, **Supplementary Table 2** and **3**). We identified 27 compounds that did not induce cell death on their own, but enhanced cell death only after transient caspase activation, with varying potencies (**Table 2**). Of these, 7 inhibited anastasis at all concentrations tested (Cdk1/2 inhibitor III, PF-3758309, BMS-754807, PHA-848125, Dinaciclib, MGCD-265, Lestaurtinib; **Supplementary Figure 1**, **Table 2**). We conclude that some kinase inhibitors promote cell death generally whereas others specifically enhance cell death only after direct, sublethal caspase activation.

**Table 2.** Kinase inhibitors reducing survival only in combination with transient caspase activation.

| | 3.3μM | 1.1μM | 0.37μM | 0.12μM | 0.04μM | 0.01μM |
| --- | --- | --- | --- | --- | --- | --- |
| JNJ-7706621 | ✓ | x | x | x | x | x |
| CHEMBL2336015 | ✓ | x | x | x | x | x |
| JAK3 inhibitor VI | ✓ | ✓ | x | x | x | x |
| NCGC00244250-01 | ✓ | ✓ | x | x | x | x |
| PD-173955 Analogue 1 | ✓ | ✓ | x | x | x | x |
| Tivozanib | ✓ | ✓ | x | x | x | x |
| Roscovitine | ✓ | ✓ | x | x | x | x |
| PKC412 | ✓ | ✓ | ✓ | x | x | x |
| AMG-51 | ✓ | ✓ | ✓ | x | x | x |
| Go 6976 | ✓ | ✓ | ✓ | x | x | x |
| PRT-060318 | ✓ | ✓ | x | x | x | ✓ |
| Motesanib | x | x | ✓ | ✓ | ✓ | x |
| BIIB-057 | ✓ | ✓ | x | x | ✓ | x |
| LDN 193189 | ✓ | ✓ | ✓ | x | x | x |
| Neratinib | ✓ | ✓ | ✓ | ✓ | x | x |
| Crenolanib | ✓ | ✓ | x | ✓ | x | ✓ |
| Bosutinib | ✓ | ✓ | x | ✓ | ✓ | ✓ |
| DCC-2036 | ✓ | ✓ | ✓ | ✓ | ✓ | x |
| NCGC00348110 | ✓ | x | ✓ | ✓ | ✓ | ✓ |
| Regorafenib | ✓ | ✓ | ✓ | x | ✓ | ✓ |
| Cdk1/2 inhibitor III | ✓ | ✓ | ✓ | ✓ | ✓ | ✓ |
| PF-3758309 | ✓ | ✓ | ✓ | ✓ | ✓ | ✓ |
| BMS-754807 | ✓ | ✓ | ✓ | ✓ | ✓ | ✓ |
| PHA-848125 | ✓ | ✓ | ✓ | ✓ | ✓ | ✓ |
| Dinaciclib | ✓ | ✓ | ✓ | ✓ | ✓ | ✓ |
| MGCD-265 | ✓ | ✓ | ✓ | ✓ | ✓ | ✓ |
| Lestaurtinib | ✓ | ✓ | ✓ | ✓ | ✓ | ✓ |

**Table 2**. Compound concentrations inhibiting anastasis (✓) are shown in red (see **Methods Section 4.5** and **Methods Section 4.6**). All other concentrations are labeled with an x.

### 3. Kinome regularization implicates multiple pathways in anastasis

To determine which individual kinases might play a role in the regulation of cellular resilience, we ran a kinome regularization analysis, which aims to explain the anastasis phenotype (the percentage of cells that survive caspase activation) as a function of the combined activity of a minimal number of kinases (39,40) (**Methods Section 4.10**) (**Figures 1A-B**). Using the previously compiled inhibitor-kinase interaction matrix (39,40) and correlating it with the variation in anastasis observed for each treatment (**Figure 4**), we applied an analytical framework similar to the one described by (39,40) (**Methods Section 4.10**). This analysis identified 46 kinases as putative regulators of cell survival following direct caspase activation (**Table 3**). Of these, Rho kinase (ROCK) is a well-established inhibitor of cell survival in some cell types (41–46), which nevertheless promotes survival of others (47–51), and can have different effects according to the duration of treatment or the adhesion status (50). Additional cytoskeletal and adhesion regulators, including protein tyrosine kinase 2 beta (PTK2B/FAK2), p21-Activated kinases (PAK1, PAK2 and PAK3), myosin light chain kinase (MYLK), Discoidin domain receptor tyrosine kinase 2 (DDR2), and CDC42 binding protein kinase β and ɣ (CDC42BPB, CDC42BPG) also emerged as candidates. These findings were intriguing because cell rounding is a defining characteristic of apoptotic cells while spreading and re-attachment are hallmarks of cells that recover in this experimental setup. We also identified several MAP kinases (MAP3K3, MAP3K7, MAP3K8, MAP3K19, MAP3K20/ZAK, MAP2K5, MAPK9/JNK2, MAPK10/JNK3), and a variety of kinases involved in growth factor signaling (FGFR1, FGFR3, FGFR4, INSR, SGK3, TEK/TIE-2, FER). In addition, we identified inhibitor of nuclear factor kappa B kinase subunit beta (IKBKB), a kinase of the NF-κB inflammatory pathway, which was recently shown to promote anastasis in colorectal cancer cells (21). These results suggested that some, but not all pathways that promote cell survival generally also specifically enhance survival of HeLa iCasp3-GC3AI cells exposed to direct caspase activity.

**Figure 4.**
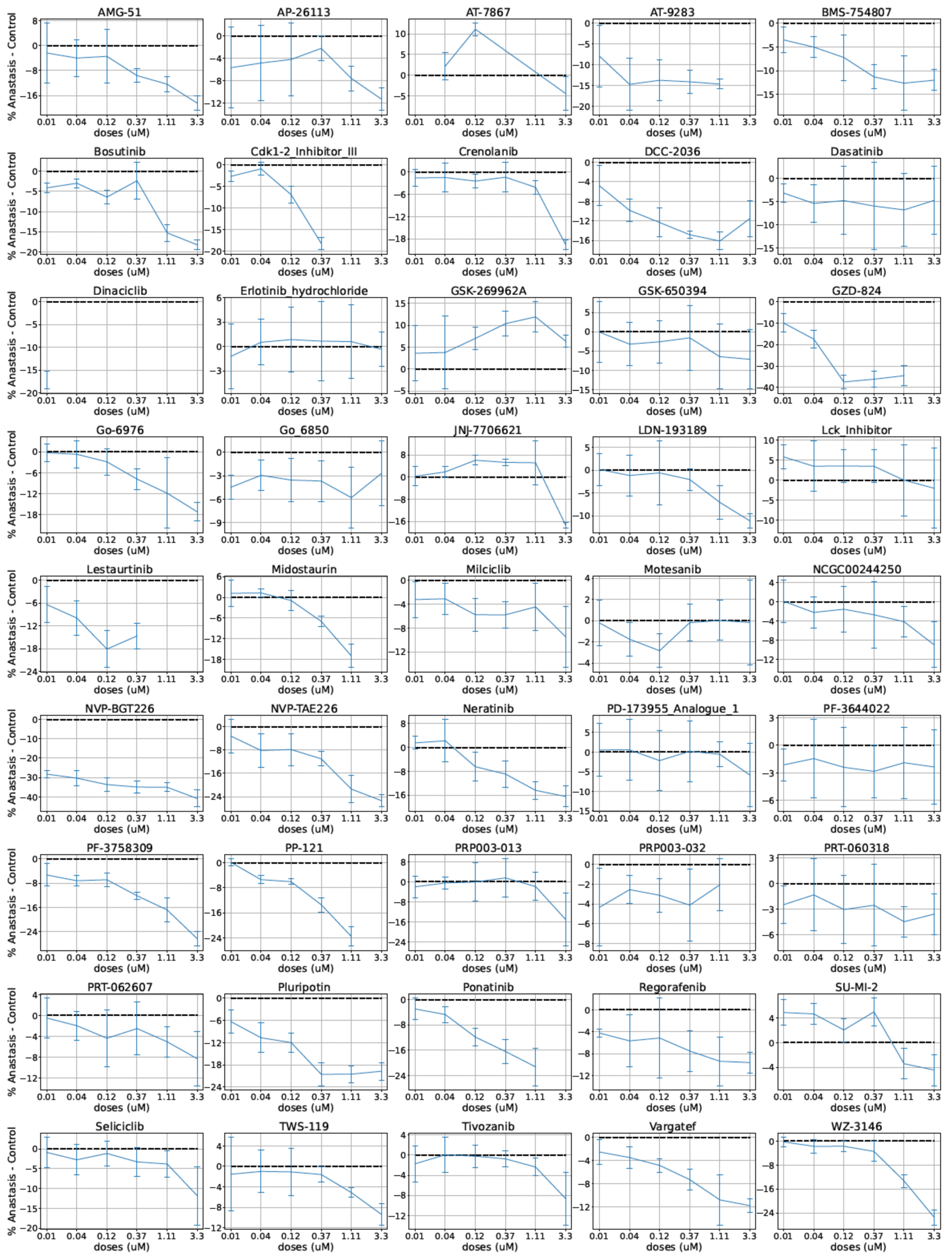
**Anastasis** phenotypes as a function of concentration for each inhibitor tested.

**Table 3.** Kinome regularization hits in HeLa iCasp3-GC3AI cells.

| Kinase | Weight | Kinase | Weight |
| --- | --- | --- | --- |
| IKBKB | 0.79 | INSR | 0.11 |
| MAP3K7 | 0.66 | INSRR | 0.1 |
| MATK | 0.63 | MAPK10/JNK3 | 0.09 |
| EPHA7 | 0.58 | STK25 | 0.09 |
| PTK2B/FAK2 | 0.54 | MYLK | 0.08 |
| FGFR4 | 0.36 | BRAF | 0.07 |
| FES | 0.33 | GRK1 | 0.07 |
| PAK2 | 0.3 | PLK1 | 0.07 |
| HIPK4 | 0.29 | MAP3K3 | 0.06 |
| PAK3 | 0.29 | FER | 0.05 |
| MAPK9/JNK2 | 0.28 | NEK4 | 0.05 |
| MAP3K19 | 0.27 | SRPK3 | 0.03 |
| MAP3K20/ZAK | 0.26 | TTBK1 | 0.03 |
| ULK3 | 0.25 | DDR2 | 0.03 |
| EPHB3 | 0.24 | STK26 | 0.02 |
| TESK2 | 0.21 | PLK2 | 0.02 |
| FGFR3 | 0.18 | MAP3K8 | 0.01 |
| MELK | 0.16 | VRK2 | 0.01 |
| PAK1 | 0.16 | SGK3 | -0.1 |
| SIK1 | 0.14 | CDC42BPB | -0.15 |
| FGFR1 | 0.13 | CDC42BPG | -0.17 |
| TEK/TIE2 | 0.12 | ROCK1 | -0.38 |
| MAP2K5 | 0.11 | ROCK2 | -0.5 |

**Table 3**. Predicted relative contributions (weights) of kinases identified as putative regulators of anastasis. A positive kinase weight (blue shading) predicts that the kinase promotes anastasis whereas a negative kinase weight (red shading) suggests that the kinase impairs anastasis.

### 4. ROCK inhibition promotes anastasis of HeLa iCasp3-GC3AI cells

Given the implication of cell adhesion and growth factor signaling, we tested whether targeting these key cellular processes could influence cell survival following direct caspase activation. The Rho-associated protein kinase ROCK regulates cell proliferation, adhesion, migration, invasion, survival and death through its effects on actomyosin contractility. Although ROCK inhibition promotes survival of stem cells (41–46), ROCK is required for survival of other cell types (47–51), and has been reported to inhibit proliferation in HeLa cells (52). Therefore, it was striking that 41nM of the ROCK inhibitor chroman I (41) induced a small but significant increase in anastasis in HeLa iCasp3-GC3AI cells with medium to high caspase activity (**Figure 5**).

**Figure 5.**
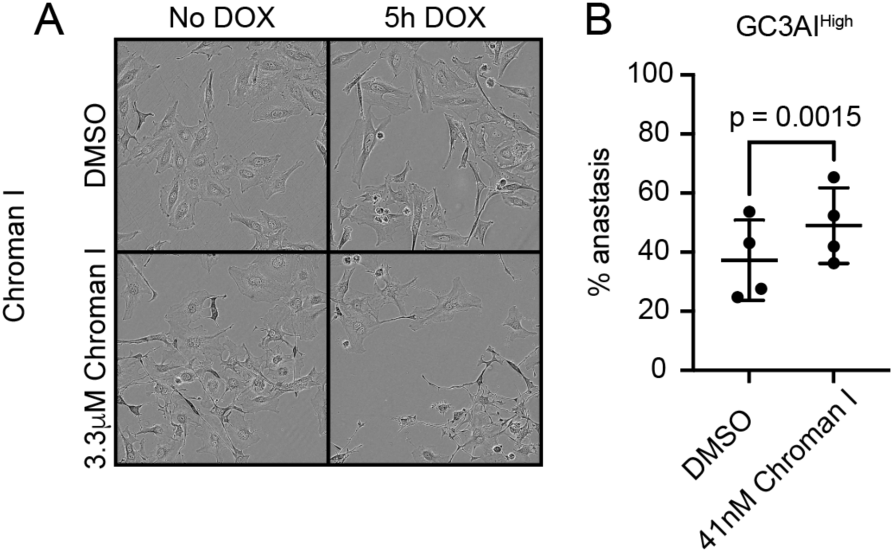
Chroman I promotes anastasis. **A.** Cells grown in complete media 0.1% DMSO (control) or complete media + 3.3µM Chroman I at the end of the experiment. At high concentrations, Chroman I affected cell morphology, prompting us to quantify anastasis manually. **B.** Dot plot illustrating the percentage of anastasis in cells with medium to high levels of GC3AI⁺ signal (detected by the Incucyte threshold mask). Error bars show the mean ± SD from n=4 independent experiments. Statistical analysis: paired t-test.

We next inhibited the cytoskeletal regulators Rho, Rac, Cdc42, PTK2B, myosin, myosin light-chain kinase, and the EMT regulator ZAK (53). We also tested whether cell adhesion could influence survival after direct caspase activation by perturbing integrin signaling and focal adhesion dynamics (see **Methods Section 6**). Unsurprisingly, at high concentrations, many of the compounds tested exhibited toxicity, even in the absence of caspase activation (**Supplementary Figures 4-5**). For example, the Rac/Cdc42 inhibitor MBQ-167 caused mitotic arrest on its own as well as a small reduction in anastasis. The Rac inhibitor NSC23766 did not, suggesting that inhibition of Rac and Cdc42 might be necessary to achieve a detectable effect. At lower concentrations, most inhibitors had limited effects on survival of HeLa iCasp-GC3AI cells with activated caspase (**Supplementary Figures 4-5**). Rac and Cdc42 activate the P21-activated kinases (PAKs). The PAK inhibitor G-5555 caused a small reduction in anastasis. However, another PAK inhibitor (FRAX-597) did not. The effects of combining selective inhibitors were also modest (**Supplementary Figure 5O-P**). Overall, these results suggest that anastasis can be enhanced or inhibited by pleiotropic kinases, but there might be substantial redundancy amongst individual kinases because the only individual kinase that caused a measurable effect on anastasis in these conditions was ROCK. The strong representation of cell adhesion and growth factor signaling pathways prompted us to test whether extracellular cues such as substrate composition, cell–matrix adhesion, or serum constituents might influence recovery from caspase activation.

### 5. Growth factor signaling promotes survival downstream of direct effector caspase activation

Among the targets identified by the kinome regularization analysis (**Table 3**), growth factor signaling emerged as a candidate pathway. Growth factors have long been known to promote cell growth and survival (54–56). To address their contribution to HeLa iCasp3-GC3AI survival after direct effector caspase activation, we challenged cells with DOX for 5h and then washed them with complete media (+10% Fetal Bovine Serum, FBS), FBS-free media, or FBS-free media supplemented with combinations of growth factors at high concentrations (**Figure 6**). Washing cells with media containing 10% FBS as well as combinations of multiple growth factors significantly increased anastasis compared to FBS-free media (**Figure 6**, **Supplementary Figures 6-7**). High concentrations of insulin, and to a lesser and more variable extent IGF2, could also support anastasis on their own (**Supplementary Figures 6-7**). These results indicate that the same signaling traditionally known to prevent apoptosis can modulate the response to direct caspase activation and that combinations of factors, especially whole serum, were more effective than individual factors.

**Figure 6.**
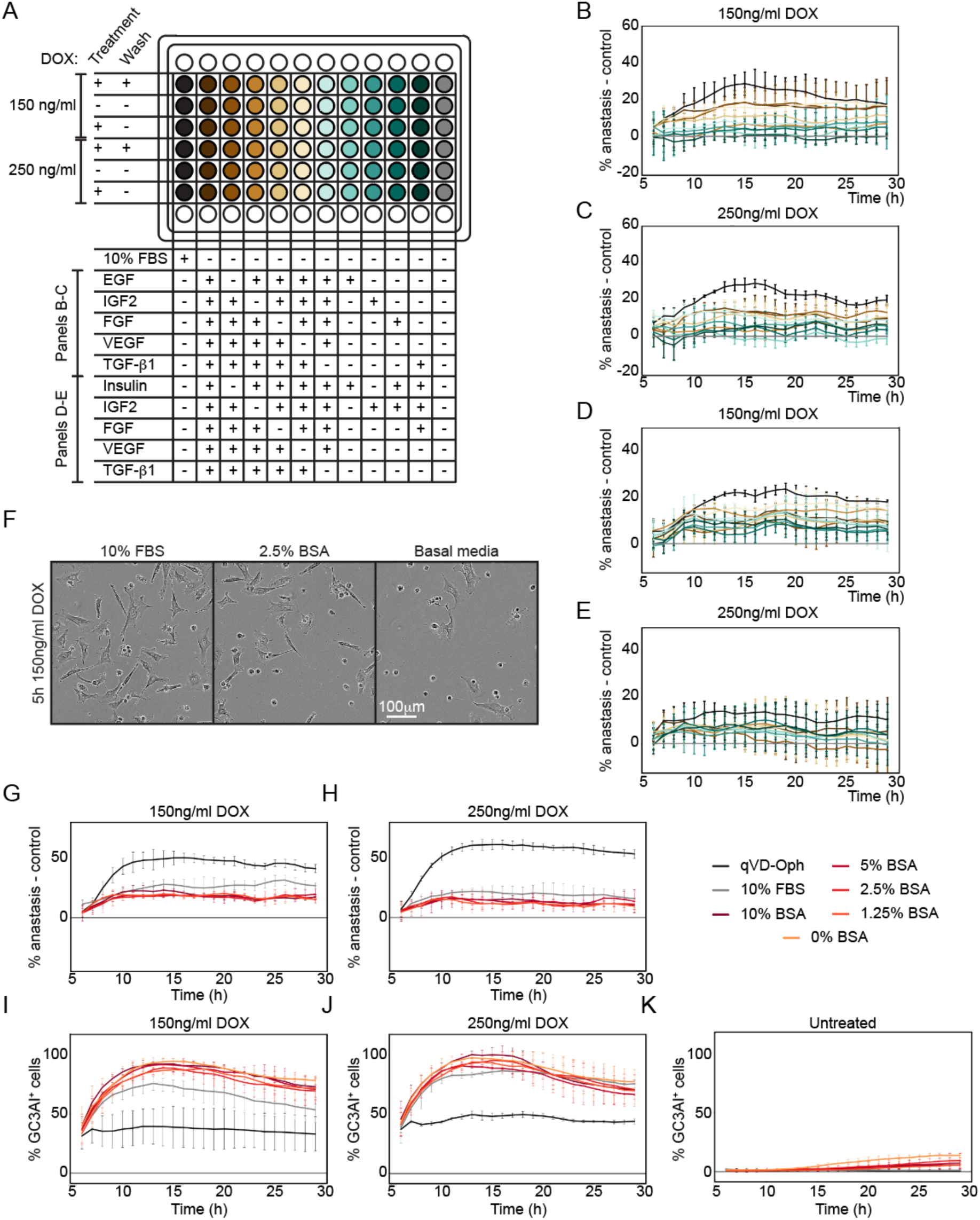
Growth factor signaling supports recovery from direct effector caspase activation. **A.** Experimental design. iCasp3-GC3AI cells were exposed transiently or continuously to 150ng/ml or 250ng/ml DOX or vehicle control. After 5h in complete media, cells were washed with complete media (10% FBS), FBS-free media, or FBS-free media with growth factors, alone or in combination. **B-E.** Percentage of cells undergoing anastasis relative to the control (base media, FBS-free) for cells exposed 5h to 150ng/ml DOX (B, D) or 250ng/ml DOX (C, E). Data were normalized by subtraction. Different treatments are shown in distinct colors, as outlined in the plate map (A). Data and error bars were smoothed. Error bars represent the mean ± SD from n=2 independent experiments. Individual treatments are shown in **Supplementary Figures 6-7**. **F-H.** BSA treatment promotes anastasis. **F.** Comparison of effects of 10% FBS, 2.5% BSA, and basal media in cells transiently exposed to 150ng/ml DOX on cell morphology at the end of the experiment. **G-H.** % anastasis normalized to the control (basal media) for cells exposed 5h to 150ng/ml DOX (G) or 250ng/ml DOX (H). 20% BSA was lethal (not shown). **I-K.** % GC3AI^+^ cells for cells exposed 5h to 150ng/ml DOX (I), 250ng/ml DOX (J), and for cells that were not exposed to DOX (K). Different treatments are shown in distinct colors. Data and error bars were smoothed. Error bars represent the mean ± SD from n=2 independent experiments.

FBS is routinely used in cell culture to promote cell survival and proliferation. Besides being a rich source of growth factors, FBS also provides hormones and nutrients. Albumin is the major protein in serum, which binds and transports amino acids, lipids, and other factors, and it has been shown to have anti-oxidant properties (57) and to carry lipids that can provide protection against apoptotic stimuli (58,59). Therefore, we tested whether Bovine Serum Albumin (BSA) alone could support cell survival from direct caspase activation. We transiently exposed HeLa iCasp3-GC3AI cells to DOX and washed with media containing different concentrations of BSA. We used heat-fractionated albumin, which is essentially free of growth factors and other heat-labile proteins. We found that BSA alone exerted a protective effect, though less than that of serum (**Figure 6**). We conclude that heat-labile growth factors and BSA contribute to the anastasis-promoting activity of FBS.

In contrast to serum and BSA, we were unable to detect effects of fibronectin, collagen I or matrigel (laminin-rich) on anastasis. No change in the fraction of surviving iCasp3-GC3AI cells could be detected, although cell morphology was clearly affected, indicating that the treatments were effective (**Supplementary Figure 8**). Fibronectin coating induced iCasp3-GC3AI cell spreading and reduced the % of cells that activated GC3AI^+^ before DOX addition (**Supplementary Figure 8C-E**), presumably by preventing caspase activation, consistent with the established role for integrin signaling in promoting cell survival generally (60–64). The lack of effect on anastasis demonstrates that not all signals that promote cell survival generally also enhance iCasp3-GC3AI anastasis.

The findings that serum promotes iCasp3-GC3AI anastasis whereas extracellular matrix molecules do not may reflect the fact that HeLa cells generally, and iCasp3-GC3AI cells in particular, are maintained in the presence of serum and in the absence of fibronectin, collagen, or matrigel, and are thus likely adapted to, or even addicted to, those conditions.

### 6. Akt signaling integrates survival stimuli to sustain anastasis

The kinase Akt is a central hub which converts growth factor signaling into pro-survival stimuli (58,65,66). Akt directly phosphorylates and inhibits pro-apoptotic proteins (27,30,67,68), and even converts them into anti-apoptotic effectors (36). Akt also promotes anastasis in *Drosophila* wing discs (14). Although Akt did not appear in the initial list of kinase hits, it emerged after a slight reduction in the kinase selectivity stringency (see details in **Methods Section 4.10**).

To assess whether Akt promotes survival after direct effector caspase activation in mammalian cells, we transiently exposed HeLa iCasp3-GC3AI cells to DOX and monitored recovery in the presence or absence of the selective inhibitor ALM301 (69,70). We found that Akt inhibition substantially prevented survival (**Figure 7A-B**). Interestingly, in the absence of caspase activation (-DOX), ALM301 only induced cell death at the two highest concentrations tested (**Figure 7C**), suggesting that cells exposed to caspase activation become selectively more reliant on Akt signaling than unexposed cells for survival.

**Figure 7.**
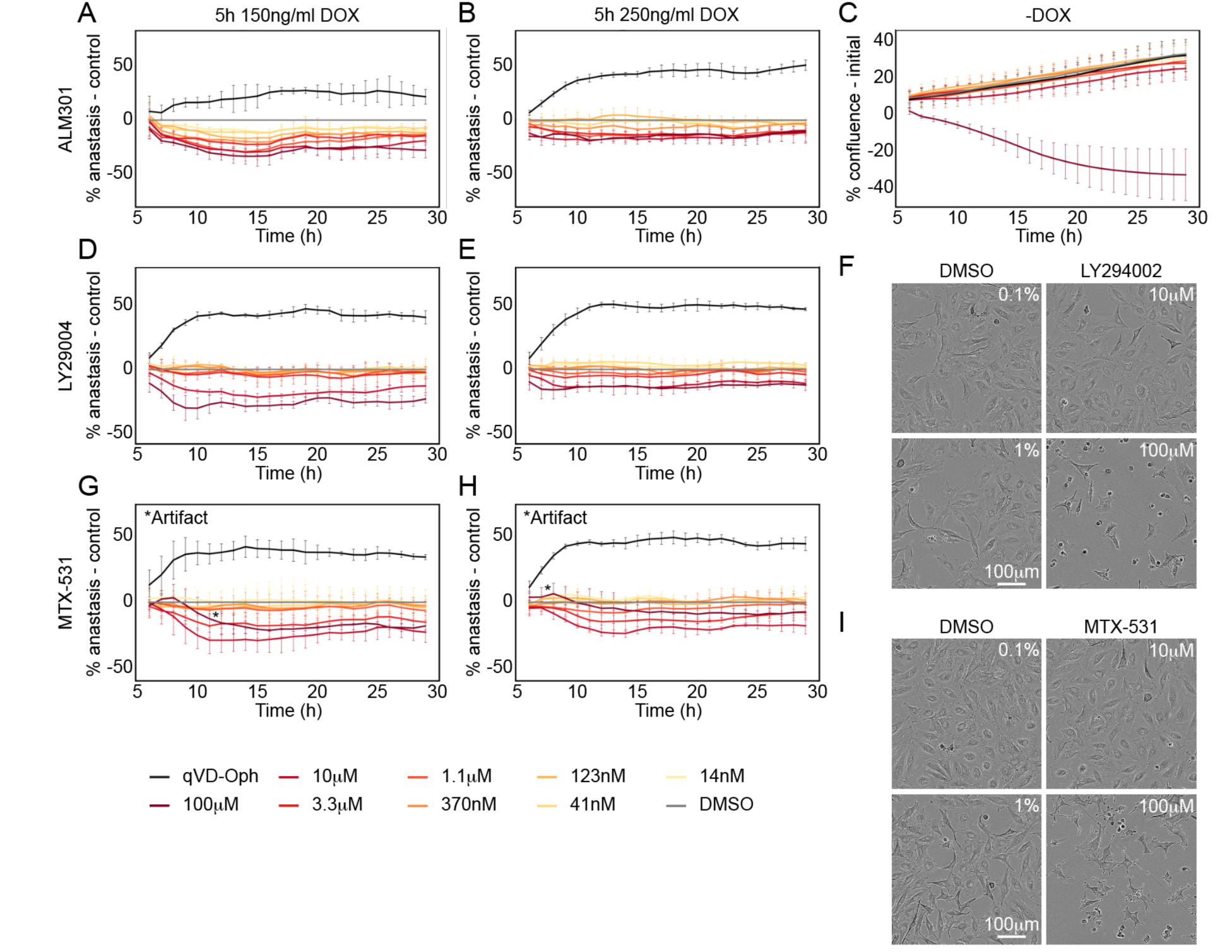
Effect of Akt inhibition on cell survival with and without caspase activation. **A-B, D-E, G-H.** Plots showing the percentage of anastasis normalized to the control in cells treated 5h with 150ng/ml (A, D, G) or 250ng/ml DOX (B, E, H), and washed with different concentrations of the Akt inhibitor ALM301 (A, B), the PI3K inhibitor LY29004 (D, E), or the dual EGFR/PI3K inhibitor MTX-531 (G, H). Although 100µM MTX-531 is the most lethal condition, the plotted curve does not fully capture this effect because extensive cell death led to aggregation, which interfered with accurate detection of anastatic cells and artificially increased their measured frequency (G-H, *Artifact). Different treatments are shown in distinct colors. The positive control (qVD-Oph) is shown in black. Data were normalized by subtraction. Data and error bars were smoothed. Error bars represent the mean ± SD from n=2 independent experiments. **C.** Plot showing the percentage confluence normalized to its initial value in cells that were not exposed to DOX for ALM301 treatment. **F, I.** Images comparing the effects of 10µM and 100µM LY294002 (F) or MTX-531 (I) and the corresponding DMSO controls in cells that were not exposed to DOX at the end of our experiment.

Phosphoinositide-3-kinase (PI3K) activates Akt (71–73). Supporting the idea that PI3K/Akt signaling contributes to anastasis, treatment with the PI3K inhibitor LY294002 diminished recovery after transient caspase activation (**Figure 7D-E**). LY294002 at 10 µM reduced survival without inducing apoptosis in no DOX controls within the timeframe of our experiment (**Figure 7F**). LY294002 is typically used in the micromolar range, so we cannot rule out that 10 µM might have one or more off-target effects. The observation that the dual EGFR/PI3K inhibitor MTX-531 (74) had similar effects supports the potential role of PI3K in survival following caspase activation.

## Discussion

Cell recovery following sublethal caspase activation, or anastasis, contributes to tissue repair after extreme injury but can also promote cancer cell survival from chemotherapy. Therefore, elucidating the underlying mechanisms and identifying pharmacological regulators of the process is important. To investigate the mechanisms and uncover kinase inhibitors that regulate survival after direct effector caspase activation, we conducted an unbiased screen beginning with pleiotropic inhibitors in HeLa iCasp3-GC3AI cells. We identified multiple pleiotropic kinase inhibitors that enhanced or inhibited anastasis at concentrations that did not affect cell survival in the absence of sublethal caspase activation.

Kinome regularization identified individual kinases that might regulate anastasis. Among the hits were 9 cytoskeletal and adhesion regulators, 8 MAP kinases, and 7 kinases involved in growth factor signaling. Our results suggest that these may have overlapping or redundant functions because, with the exception of ROCK and Akt, pleiotropic inhibitors were more effective than specific inhibitors. ROCK functions primarily downstream of the small GTPase Rho, which regulates actomyosin contractility. While relatively low levels of Rho/ROCK signaling promote adhesion to extracellular matrix and focal adhesion formation, high levels of Rho/ROCK can cause cell rounding and detachment, which are defining features of apoptotic cells generally and DOX-treated HeLa iCasp3-GC3AI cells specifically. This dual function of ROCK in promoting attachment and detachment is consistent with the possibility that a low concentration of ROCK inhibitor promotes anastasis by lowering the level of ROCK activity just enough to favor attachment rather than detachment. Such dual functions might represent a general challenge for identifying regulators of anastasis, in addition to functional overlap or redundancy. This dual function of ROCK may also underlie its varied and context-dependent effects on cell survival in diverse cell types and conditions, as distinct cell types may have and/or require different levels of ROCK activity depending on their cell state and microenvironment.

We found that growth factors and BSA promote HeLa Casp3-GC3AI cell survival after caspase activation. Supporting the notion that simultaneous activation of multiple pathways is required, FBS was more effective than individual growth factors in promoting survival. Our findings demonstrate that pro-survival signaling not only prevents caspase activation, but also provides cells the opportunity to recover after effector caspases have been activated.

Importantly, we identify Akt as a central signaling node for survival, extending observations previously made in *Drosophila* (14). Consistent with Akt’s role in preventing apoptosis (27,30,36,67,68) our work positions Akt-dependent signaling as a critical determinant of whether a cell dies or recovers following caspase activation.

HeLa cells are typically cultured in the presence of serum but the absence of adhesive coatings, which may explain why neither fibronectin, collagen, nor laminin-rich matrigel measurably affected anastasis of HeLa iCasp3-GC3AI cells. The idea emerges then that cells growing in distinct microenvironments, for example at the edges of solid tumors where stiff extracellular matrix is abundant, versus other locations where for example, proximity to blood vessels might provide increased growth factor signaling, might adapt to those locations and acquire specific molecular dependencies and vulnerabilities. Thus, another form of tumor heterogeneity, in addition to genetics, may be the specific factors that promote survival following caspase activation.

Beyond identifying specific mechanisms and pharmacological inhibitors, our work suggests that anastasis may represent a tunable cellular property. In regenerative contexts, transient ROCK inhibition or growth factor supplementation could enhance recovery of stressed cardiomyocytes or neurons. Conversely, in oncology, combining chemotherapy with Akt/ROCK modulators might prevent the emergence of persistent, drug-tolerant cells that drive relapse. The kinome signature we identified provides a roadmap for such targeted interventions.

## Authors contributions

M.N. and D.J.M. designed research; M.N., J.H., and G.L. performed research; M.N., J.H., G.L. and A.K. analyzed data; M.N. and D.J.M. wrote the paper; D.J.M. and L.P. supervised the work.

## Acknowledgements

We acknowledge Thomas Craig for providing the inhibitors. Maria Malafei, Felipe Da Costa Lima, and Sam Chen provided technical help. LP was supported by the NIH award AG073341. The authors acknowledge the use of the Sony MA900 activated cell sorter in the Biological Nanostructures Laboratory within the California NanoSystems Institute, supported by the University of California, Santa Barbara and the University of California, Office of the President. All plasmids were obtained from Addgene; the plasmid pCW57-MCS1-2A-MCS2 was a gift from Adam Karpf, pCDNA3-FlipGFP(Casp3 cleavage seq) T2A mCherry was a gift from Xiaokun Shu, and pCMV-dvpr-dR8.2 and pCMV-VSV-G were gifts from Bob Weinberg.

## Methods

### 1. Cell Culture and Maintenance

iCasp3-GC3AI were described in (24) (CaspaseLOV cells). Briefly, HeLa cells were first engineered to stably express the caspase activity reporter GC3AI (26) by lentiviral transduction, followed by CRISPR–Cas9 knockout of endogenous caspase-3. The cells were subsequently transduced with a DOX-inducible, partially active caspaseLOV construct (25), whose activity is further enhanced by blue-light illumination. Monoclonal populations were isolated and validated after each genetic modification. GC3AI is activated upon cleavage by effector caspases, while DOX induction of caspaseLOV triggers GC3AI activation.

Cells were grown in DMEM (cat#10-569-044, Gibco by Thermo Fisher Scientific, Grand Island, NY, USA) with 10% heat-inactivated fetal bovine serum (cat#F4135, Millipore Sigma, Burlington, MA, USA; or cat#25-514H, Genesee Scientific, El Cajon, CA, USA), and 1 μg/ml puromycin (cat#A1113803, Gibco) as a selection agent. Cells were maintained at 37°C with 5% CO_2_ and 90% humidity. For experiments investigating the contribution of FBS to cell survival after caspase activation, we used product #25-514H from Genesee Scientific (lot#30224001). For experiments investigating the contribution of BSA to cell survival after caspase activation, we used product #A1470 from Millipore Sigma (Lot#0000482631, Source#0000474087).

### 2. Plate coating

We investigated the contribution of substrate composition and stiffness to cell survival after caspase activation utilizing 96 well glass bottom plates with high performance #1.5 cover glass (cat#P96-1.5H-N, Cellvis Scientific, Mountain View, CA, USA). We performed coating overnight at 37°C, and briefly rinsed the wells with room temperature PBS (cat#10010023, Thermo Fisher Scientific, Waltham, MA, USA) before plating cells. We diluted fibronectin (cat#1918-FN, R&D systems, Minneapolis, MN, USA) to 0.01mg/ml in PBS and added 100µl to each well. We diluted rat tail collagen (cat#CB354249, Thermo Fisher Scientific) to 0.08-0.11 mg/ml in ice-cold water + 0.02M acetic acid and added 100µl to each well. Growth Factor Reduced Matrigel (cat#356231, lot #1319010, Corning Incorporated, Corning, NY, USA) was thawed on ice, diluted to 100 µg/ml in ice-cold PBS and distributed at 100 µl/well.

### 3. Generation of DOX-inducible mCherry HeLa cells

mCherry HeLa cell lines were generated by lentiviral transduction. Lentiviral particles were produced in Lenti-X 293T cells (cat#632180, Takara Bio USA, San Jose, CA, USA) using a transfer vector (pCW57 DOX-inducible vector; cat#71782, Addgene) modified to express mCherry, the envelope plasmid pCMV-VSVG (cat#8454, Addgene), and the packaging vector pCMV-dvpr-dR8.2 (cat#8455, Addgene). Detailed cloning and lentiviral particles production are described in the **Supplementary Methods**.

### 4. Kinome regularization

#### 4.1 Pleiotropic kinase inhibitors

As described in (39,40), the full set of kinase inhibitors to use in Kinome Regularization was selected from hundreds of inhibitors, previously characterized by applying Principal Components analysis and selecting the drugs that are minimally redundant and maximally diverse. Among the original 58 Principal Compounds, 48 contained a dose that significantly inhibited (<0.5 kinase residual activity) at least 10% of the kinome. So, we excluded the remaining 10 compounds from our analysis as well as NCGC00344999 because it was not available. For Cdk1/2 Inhibitor III, we used compound CAS#443798-47-8 instead of CAS#443798-55-8.

Most inhibitors screened are listed in **Supplementary Table 1**. We also purchased the following compounds: Cdk1/2 Inhibitor III (CAS: 443798-47-8, cat#18859, Cayman Chemicals, Ann Arbor, MI, USA), JAK3 Inhibitor VI (CAS: 856436-16-3, cat#42-012-65MG, Thermo Fisher Scientific,), Lestautinib (CAS: 111358-88-4, cat#12094, Cayman chemicals), PKC 412 (CAS: 120685-11-2, cat#10459, Cayman chemicals), RK 2446 (CAS: 213743-31-8, cat#15135, Cayman chemicals), Vargatef (CAS: 928326-83-4, cat#V2099.5, United States Biological Corporation, Salem, MA, USA).

#### 4.2 Kinome regularization: inhibitors screening

Cells were seeded at 50-80% confluency in 96 well plates (cat#25-109, Genesee Scientific) in complete DMEM media. All DOX and drug treatments were made using DMEM, 15% FBS (cat#F4135, lot #20H250, Millipore Sigma) + 1µg/ml puromycin. Cells were treated as shown in **Figure 1D-E**. After 5h of treatment with media or media + DOX, cells were washed with media with or without DOX containing a given concentration of kinase inhibitor, a positive control (qVD-Oph, cat#A1901, APExBIO, Houston, TX, USA), or a mock treatment (DMSO, cat#D45-40, Millipore Sigma). qVD-Oph and DOX concentration were adjusted as described in the **Supplementary Methods**. For all compounds we ran n=2 independent experiments, except for NCGC00244250-01, for which the two biological replicates were run in parallel.

#### 4.3 Kinome regularization: IncuCyte imaging and image analysis

Cells were imaged 30min pre-treatment, 30min post-treatment, and then hourly for at least 30h using an IncuCyte Zoom system (Sartorius, Göttingen, Germany) equipped with a Dual Color Model 4459 filter module and a Nikon 20x objective (Green Acquisition Time=400msec), or an IncuCyte S3/SX1 using the 20X objective with G/R Optical Module (Green Acquisition Time=300msec) and the Adherent Cell-by-cell scan type. To identify cells activating caspase, we generated a mask for GC3AI as described in the **Supplementary Methods**, using a threshold that maximized detection of GC3AI-positive cells in DOX-treated wells while minimizing detection in untreated wells.

#### 4.4 Kinome regularization: IncuCyte data handling

For each treatment and time point, we determined the fraction of GC3AI⁺ cells that remained alive after caspase activation (% anastasis). Then, we calculated the difference between each condition and the mock. Data were averaged across replicates.

This method is intentionally insensitive to compounds that alter the total number of cells (GC3AI^+^ + GC3AI^-^) or the total number of GC3AI^+^ cells without altering the fraction of GC3AI^+^ cells that die. This can occur for compounds affecting cell cycle progression or inducing more or less penetrant/detectable GC3AI activation above the threshold used to determine GC3AI positivity. Compounds that increase GC3AI activation are also likely to cause an increase in the fraction of dying cells at the population level, which might not be detected. Compounds that decrease GC3AI activation are also likely to cause an increase in the fraction of surviving cells at the population level, which might not be detected.

Detailed description of data processing for visualization and handling is reported in the **Supplementary Methods**.

#### 4.5 Kinome regularization: analysis of inhibitor effects on anastasis

To evaluate the effect of each treatment on anastasis, we examined how it changed the fraction of living GC3AI^+^ cells in samples transiently (5h) exposed to DOX. We processed the data as described in **Methods Section 4.4** and corrected them as described in **Methods Section 4.8** and **4.9**. To classify the overall effect on anastasis as negative, neutral, or positive (**Table 1**, **Table 2**, and **Supplementary Table 2**), we assigned a score to each treatment as described in the **Supplementary Methods**.

#### 4.6 Kinome regularization: analysis of compound toxicity in the absence of caspase activation

To determine whether any compound increased caspase activation (GC3AI) in the absence of DOX (**Supplementary Figure 2**), we used a numerical threshold that best captured visually detectable increases in GC3AI. The threshold was determined as described in the **Supplementary Methods**. Results are summarized in **Supplementary Table 3**.

#### 4.7 Kinome regularization: analysis of compounds interference with DOX induction

To determine whether kinase inhibitors interfered with DOX-induced caspase expression, we employed multiple complementary approaches: (1) we assessed whether anastasis occurred in samples continuously exposed to DOX (**Supplementary Figure 1C**) (**Methods Section 4.7.1**), (2) we analyzed GC3AI induction in samples treated with DOX for 5h followed by a wash (**Supplementary Figure 9**) (**Methods Section 4.7.2**), and (3) we examined a subset of compounds for their ability to disrupt DOX-inducible mCherry expression by western blot (**Methods Section 4.7.3**) (**Supplementary Figure 10**). We believe these approaches reliably identified kinase inhibitors that affect DOX-inducible gene expression; however, if such interference did not translate into increased survival in unwashed samples or enhanced anastasis, it might not have been detected.

##### 4.7.1 Detection of anastasis in samples continuously exposed to DOX

In this experimental setup, most cells continuously exposed to DOX died by 24h. To identify compounds that might interfere with DOX induction we searched for treatments that increased the fraction of living GC3AI^+^ cells under continuous DOX exposure (**Supplementary Table 4**) as described in the **Supplementary Methods**.

##### 4.7.2 Detection of GC3AI induction in samples treated with DOX for 5h and washed

To further evaluate how kinase inhibitors affected DOX induction and caspase activation, we quantified the number of GC3AI⁺ cells for each treatment and normalized these values to the DMSO control (GC3AI⁺ inhibitor/GC3AI⁺ DMSO)(**Supplementary Figure 9**). The effect of each compound was determined as described in the **Supplementary Methods** and is summarized in **Supplementary Table 5**.

##### 4.7.3 Western blot

To test whether compounds enhance survival after caspase activation by reducing DOX-inducible gene expression (transcription or translation) or by reducing caspase activity, we used HeLa cells encoding DOX-inducible mCherry (see **Methods Section 3**). The compounds to test were selected because they promoted the survival of cells transiently exposed to DOX (RK-24466), and/or because they inhibited cell death in DOX-treated, unwashed samples (all others) (**Supplementary Table 6**). The transcriptional inhibitor Actinomycin D was used as a positive control (100nM), and caused high levels of cell death. To facilitate detection of perturbations in mCherry expression, inhibitors were added together with DOX and were not washed (**Supplementary Figure 10A**). Samples were treated ∼16h with 1µg/ml DOX and a non-toxic, artifact-free concentration identified at the IncuCyte for each compound (**Supplementary Table 6**). Protein extraction and western blot were performed as described in the **Supplementary Methods**. Results are shown in **Supplementary Figure 10B-C**.

#### 4.8 Kinome regularization: strategy to normalize anastasis levels in compounds interfering with DOX induction

To normalize anastasis levels in compounds that interfere with DOX induction (**Methods Section 4.7**), we used the following strategy (**Supplementary Figure 3**): (1) we quantified red fluorescence intensity in HeLa cells expressing a DOX-inducible mCherry construct treated with DOX, with or without a given kinase inhibitor; (2) we determined how anastasis levels following DOX treatment in HeLa cells expressing caspaseLOV correlated with mCherry expression in cells carrying the DOX-inducible mCherry construct (calibration); and (3) we applied this calibration to correct anastasis values in our screening dataset. The detailed procedure is reported in **Supplementary Methods**. This approach assumes that the mechanisms affecting mCherry induction and anastasis are analogous. **Supplementary Table 7** summarizes the corrected treatments. Corrected data are presented in **Figure 2**, **3**, **4**, **Supplementary Figure 1**, **9**, **Table 1**, **2**, **Supplementary Table 2**, **5**, and **10**. Original and corrected data are reported in **Supplementary Figure 1B** and **9B**.

#### 4.9 Kinome regularization: strategy to normalize anastasis levels in compounds exhibiting artifacts

For a subset of compounds, fluorescence artifacts interfered with the reliability of the measured anastasis values. To address this, we developed an extrapolation-based strategy to predict expected anastasis levels in the affected treatments. The detailed procedure is described in the **Supplementary Methods**. Although this approach was developed for anastasis measurements, the same logic was applied to the induction data. This normalization approach was applied to the following compounds and concentrations: 3.33μM Vargatef (autofluorescent), 3.33μM Ponatinib (dead cell aggregation), and 3.33μM PF-3644022 (precipitates in green crystals). 3.33μM and 1.11μM PKI-SU11274 also presented artifacts (autofluorescent), but were corrected using the method described in **Methods Section 4.8**. Corrections are included in **Figure 2**, **4**, **Supplementary Figure 1**, **9**, **Supplementary Table 2**, **5** and **10**. Original and corrected data are reported in **Supplementary Figure 1B** and **9B**.

##### 4.9.1 Correction by Manual Counting

GZD824 treatment (3.33-0.12 µM) induced dead cell aggregation that compromised automated quantification. Anastasis was therefore manually quantified in 2 of 4 images per affected well at 13, 15, and 18 h by counting living and dead GC3AI⁺ cells detected by the Incucyte (**Methods Section 4.3**) and normalized to the matched DMSO control for each replicate. Corrections are included in **Figure 4**, **Supplementary Figure 1**, **9**, **Supplementary Table 2**, **5** and **9**. Original and corrected data are reported in **Supplementary Figure 1B** and **9B**.

#### 4.10 Kinome regularization analysis

Among the 47 screened drugs, JAK3 Inhibitor VI and MGCD-26 were not present within the inhibitor-kinase interaction matrix and were eliminated from our analysis. We also excluded NVP-BGT226, as it exhibited clear outlier behavior, producing stronger inhibition of anastasis than all other compounds even at the lowest concentrations tested. As a result, its dose–response profile fell outside the range modeled by the regression, and all doses of this compound were treated as outliers and excluded from analysis. 3.33 µM DCC-2036, Crenolanib, GSK-269962A, and JNJ-7706621 were excluded because their corresponding anastasis values appeared as clear outliers in **Figure 4**.

By restricting the inhibitor–kinase interaction matrix to the selected subset of inhibitors (rows), we obtained a 242 × 369 submatrix, hereafter denoted *X*. Each row of *X* was then paired with the measured anastasis value of the corresponding inhibitor, yielding a response vector *Y* of length 242. We applied statistical learning methods to derive the functional relationship between *X* and *Y* and determine the kinases (covariates) affecting anastasis (response). As the number of covariates (369 kinases) in our data is larger than the number of responses (242 inhibitors), we follow the sparse regularization framework of (39,40), which uses the ElasticNet regularization to select only a relatively small subset of kinases associated with the phenotype. This approach models each phenotype value *Y_i_* as a weighted combination of kinase activities *X_ij_*, *j* = *1*, …, *369*, while penalizing for too many nonzero weights in this sum. If the weight *β_j_* in front of kinase *j* is zero, it means that changing the inhibition level of this kinase alone does not affect anastasis and we can deduce that there is no association between the two. A positive weight *β_j_* means that with all other kinase activities fixed, a unit decrease in *X_ij_* (a unit increase in inhibition of kinase *j*) leads to a decrease in anastasis by *β_j_* units. Similarly, a negative weight means that a unit decrease in *X_ij_* leads to the increase in anastasis by this number of units. To estimate the unknown kinase weights from the data *X* and *Y* at hand, the ElasticNet model minimizes the following objective over *β_j_*, *j* = *1*, … , *369* and *intercept β_0_*, that is, the anastasis level an inhibitor *i* would achieve if it fully inhibited all the kinases (*X_ij_* = *0* for all *j*):

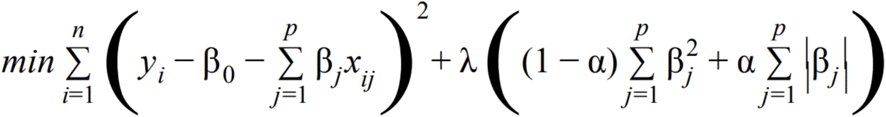

Here, *n* = *242* is the number of observations, *p* = *369* is the number of features (kinases) and (*α*, *λ*) are two penalization parameters. Intuitively, *λ* here is a complexity parameter determining the tradeoff between the number of variables chosen and the goodness of fit of the model (the higher *λ*, the less kinases are selected and the worse the model is fit); *α* is a number between 0 and 1 responsible to balance selectivity and smoothness of weights assigned to correlated features (kinases). With *α* close to 1, the model selects the least possible number of kinases by choosing no more than one kinase from a group with highly correlated inhibition profiles. On the contrary, with *α* close to 0, the model provides the largest possible set of selected kinases but assigns similar weights to highly-correlated kinases. Similarity of weights for correlated kinases is important to ensure the model identifies all the “siblings” of a candidate phenotype-associated kinase. To tune the unknown parameters *λ* and *α*, we use a Leave-One-Drug-Out Cross Validation (LODOCV) procedure. For each combination of the two parameters in a predefined grid, this method iteratively fits the model on all but one drug, predicts the phenotype for all the doses of the remaining drug, and then computes the mean absolute error between the true and predicted values. This procedure is repeated for every possible drug and concluded with measuring the average error across the *n* fits. For our data, the optimal parameters, chosen as the ones minimizing this average error, turned out to be *λ*=1 and *α*=0.55. Keeping *λ* fixed, slightly lowering *α* from its optimal value can bring out a small number of additional candidate kinases. For example, as mentioned in **Section 6**, Akt becomes a candidate kinase (non-zero weight) if we reduce *α* from 0.55 to 0.45.

### 5. Screening of selective compounds

For our candidate screening, cells were plated at low density (2.5k/well) in a tissue culture treated 96 well plate and were treated as described in **Methods Section 4.2**, including two additional concentrations (10µM and 100µM). **Supplementary Table 8** lists the inhibitors tested and their main targets. Detailed information for imaging and image analysis are available in the Supplementary Methods.

### 6. Validation of hits from kinome regularization

Small molecules were reconstituted in DMSO, and were screened and analyzed as described in **Methods Section 6**. For confluence analysis, we used the Sartorius “AI Confluence” segmentation (Adjust Size=-10px, Area>150µm^2^). We used ALM301 (CAS #1313439-71-2) to inhibit Akt (cat#HY-151504, MedChemExpress) (69), LY294002 (CAS #154447-36-6) to inhibit PI3K (cat#PHZ1144, Thermo Fisher Scientific) (75,76), MTX-531 (CAS #2791417-66-6) to inhibit EGFR/PI3K (cat#43038, Cayman Chemicals) (74), and rapamycin (CAS #53123-88-9) to inhibit mTOR (cat#HY-10219, MedChemExpress) (77).

Growth factors were reconstituted according to manufacturer instructions, and diluted in FBS-free media + 1 μg/ml puromycin. Origin and concentrations are shown in **Supplementary Table 9**. Most growth factors were used at supraphysiological concentrations.

### 7. Code availability

The code and raw data of the screening are publicly available on **GitHub**.

