## Supplementary material for "A kinome inhibitor screen implicates adhesion and growth factor signaling in cellular recovery after caspase activation"

Includes: Supplementary Methods, 11 Supplementary Figures, 10 Supplementary Tables, and Supplementary Bibliography

**Supplementary Methods**

**Relative to Methods Section 3.** We amplified mCherry from PCDNA3-FlipGFP(Casp3 cleavage seq) T2A mCherry (cat#124428, Addgene, Watertown, MA, USA) using the primers: forward (5' - tatataACGCGTatggtgagcaagggcgagg - 3') and reverse (5' - atatatGGATCCttacttgtacagctcgtccatgcc -3'). mCherry was ligated into a pCW57 DOX-inducible vector (cat#71782, Addgene) downstream the P2A site using MluI-BamHI restriction enzymes (cat#R3198, cat#R3136, New England Biolabs, Ipswich, MA, USA). The transfer vector was delivered to HeLa cells (cat#CCL-2, ATCC, Manassas, VA, USA) using lentiviral transduction. Lentivirus was packaged using the transfer vector, the envelope plasmid pCMV-VSVG (cat#8454, Addgene), and the packaging vector pCMV-dvpr-dR8.2 (cat#8455, Addgene) at a ratio of 5:1:3, respectively. Transient expression of vectors for virus production in Lenti-X 293T cells (cat#632180, Takara Bio USA, San Jose, CA, USA) was achieved using X-tremeGENE™ HP DNA Transfection Reagent (cat#6366236001, Millipore Sigma). 24hr post-transfection, cells were transferred in fresh DMEM + 10% FBS media without antibiotics. We collected viral supernatant 24hr after media change and filtered it through 0.45µm nitrocellulose syringe filter (cat#28143-352, VWR, Radnor, PA, USA) and concentrated to 10X using LentiX Concentrator (cat#631231, Takara Bio USA) according to manufacturer specifications. HeLa cells were transfected using concentrated virus via spinoculation. Briefly, 250µL of virus and polybrene (cat#TR-1003-G, Millipore Sigma) at a final concentration of 2µg/ml were added directly to 200k cells in a 6 well dish and spun 45min RT at 2000xg. The following day, cells were washed twice with PBS and media was replaced with DMEM + 10% FBS + 1µg/ml puromycin to select for successfully transfected cells. After establishment, cells were treated 24hr with 1µg/ml DOX and sorted for mCherry positivity at a Sony MA900 activated cell sorter (Sony Biotechnology, San Jose, CA, USA), expanded, and used for subsequent experiments.

**Relative to Methods Section 4.2.** qVD-Oph concentration was adjusted to obtain at least ~20% increase in survival in cells that activate GC3AI after transient DOX exposure compared to mock-treated cells (1µM to 10µM). Similarly, DOX concentration was adjusted (75ng/ml to 250ng/ml) to obtain a ~3-4 fold increase in GC3AI fluorescence by 4h30min post-treatment and ~6-8 fold increase by 5h30min post-treatment. Even so, the fraction of cells activating GC3AI was variable across experiments. Therefore, we calculated the difference between each sample and the mock for each experiment (see **Methods Section 4.4**). While qVD-Oph change in potency was most likely linked to batch variability, the variable response to DOX seemed to correlate with cell passage number and cell seeding density.

**Relative to Methods Section 4.3.** To identify cells activating caspase, we generated a mask for GC3AI. We subtracted the background using a Top-Hat method (radius=10µm, threshold=0.25GCU). We set edge sensitivity to -45, with Edge Split On, hole filling to 100µm and size adjustment to -2px. We only considered objects larger than 100µm$^2$. To distinguish between dead and live cells, we included an empirically determined eccentricity filter (living cells eccentricity >0.7). Only objects where the green signal surpassed a threshold were considered for analysis.

**Relative to Methods Section 4.4. Data visualization.** Smoothing of data was obtained using a moving average of the mean across timepoints with a window of size 3, centered around the current time point. End points (the first and last timepoints) were the average of 2 timepoints, the end and the timepoints next to it. Error bars followed the same method, applied on the standard deviation instead of the mean. **Kinome regularization.** To use as an input for the kinome regularization we averaged the unsmoothed anastasis values for time point 12, 13, 14, 15, 16, 17 and 18 because they captured substantial responses for most inhibitors and were less likely to be influenced by differences in proliferation. Corrections were made to account for compounds that interfered with DOX-inducible gene expression (see **Methods Section 4.8**) and for compounds that caused morphological or autofluorescent artifacts (see **Methods Section 4.9**).

**Relative to Methods Section 4.5.** To classify the overall effect on anastasis as negative, neutral, or positive, we assigned a score to each treatment. For each time point, we defined a scoring range as one standard deviation above and below the mean anastasis value. We then assigned a score of −1 if this range fell entirely below the control value (0), 0 if it included 0, and +1 if it lay entirely above 0. For each treatment, we next determined the mode of these scores (either across all post-wash time points, **Supplementary Table 10**; or specifically within the 12–18 hour window, **Table 1**, **Table 2**, and **Supplementary Table 2**). In cases where multiple modes occurred, their average was used.

**Relative to Methods Section 4.6.** To determine whether any compounds increased caspase activation (GC3AI) in the absence of DOX, we first examined the images to assess whether the number of $GC3AI^{+}$ cells exceeded that observed in the DMSO control at the end of our experiment. We then quantified GC3AI+ cells for each compound-only condition (-DOX) and subtracted the corresponding DMSO value. By comparing these quantitative results with the visual assessment, we identified a numerical threshold that best captured visually detectable increases in GC3AI.

To choose this threshold, we tested a range of values and, for each one, calculated: (1) how many compounds were flagged numerically but not visually, and (2) how many were flagged visually but not numerically. The threshold minimizing the sum of these mismatches was selected (cutoff=5). Any remaining discrepancies were reviewed individually. A comparison of compound toxicity with and without DOX can be made comparing Supplementary Table 2 and 3. Supplementary Table 3 summarizes treatments where GC3AI signal exceeded the threshold at ≥2 time points within the 12-18h time window.

**Relative to Methods Section 4.7.1.** To identify compounds that might interfere with DOX induction, we searched for treatments that increased the fraction of living GC3AI+ cells under continuous DOX exposure. For each treatment, we quantified anastasis (percentage of living GC3AI+ cells minus corresponding DMSO control value). Heatmap in Supplementary Figure 1C shows anastasis averaged across TP12-18. To build Supplementary Table 4, we compared these numerical results with the image-based observations to determine a cutoff value that best reflected visually detectable increases in survival. To select this cutoff, we evaluated a range of thresholds and, for each, calculated: (1) the number of treatments classified as hits by the numerical threshold but not visually, and (2) the number identified visually but not by the threshold. The threshold value that minimized the sum of these mismatches was chosen (cutoff=15). Treatments in which the percentage of living GC3AI-positive cells exceeded this threshold at more than one timepoint throughout the experiment are summarized in Supplementary Table 4.

**Relative to Methods Section 4.7.2.** To evaluate how kinase inhibitors affected DOX induction and caspase activation, we quantified the number of $GC3AI^{+}$ cells for each treatment and normalized these values to the DMSO control ($GC3AI^{+}$ inhibitor/$GC3AI^{+}$ DMSO)(Supplementary Figure 9). To further distinguish effects on viability from effects on DOX-induction, we then corrected these values to account for compound-dependent interference with DOX-inducible expression, as described in Methods Section 4.8. For each time point, we defined a scoring range as one standard deviation above and below the mean induction value. Since a compound which does not affect induction would have a mean of 1, we use 1 for the comparison. We then assigned a score of -1 if this range fell entirely below the control value (1), 0 if it included 1, and +1 if it lay entirely above 1. For each treatment, we next determined the mode of these scores within the 12-18 hour window, to classify the overall effect on induction as negative, neutral, or positive (Supplementary Table 5). In cases where multiple modes occurred, their average was used.

**Relative to Methods Section 4.7.3**. HeLa cells with DOX-inducible mCherry (Methods Section 3) were plated at a density of 200K cells/well in a six well dish (cat#3516, Corning) in DMEM + 10% FBS + 1µg/ml puromycin. The following day, cells were treated with media with or without DOX, kinase inhibitors, qVD-Oph, Actinomycin D (cat#11421, Cayman Chemicals), or DMSO

(Supplementary Figure 10). After 16h, cells were washed once with ice-cold PBS, and lysed with 100µL of RIPA Lysis Buffer (cat#89900, Thermo Fisher Scientific) supplemented with cOmplete™, Mini, EDTA-free Protease Inhibitor Cocktail and PhosSTOP™ phosphatase inhibitor cocktail tablets (cat#11836170001, cat#4906845001, Millipore Sigma). Protein concentrations were determined using Pierce BCA Assay Kit (cat#23227, Thermo Fisher Scientific) and lysates were equilibrated to the lowest sample concentration using 6X Laemmli Sample Buffer (cat#J61337.AD, Thermo Fisher Scientific), and stored at -20°C until use. After thawing on ice, samples were heated at 90°C for 10min. Proteins were separated on 4–20% Mini-PROTEAN TGX Precast Protein Gels with 15 combs (cat#4561096, Bio-Rad, Hercules, CA, USA) in 10% Tris-Glycine (cat#1610771, Bio-Rad) + 0.1% SDS (cat#1610418, Bio-Rad) running buffer. Molecular weights were determined using Precision Plus Protein All Blue Prestained Protein Standards (cat#1610373, Bio-Rad). Transfer of proteins onto nitrocellulose membrane (cat#1620115, BioRad) was carried out at 70V for 1.5h at 4°C in an aqueous buffer composed of 10% Tris-Glycine + 20% methanol (cat#A412P-4, Thermo Fisher Scientific). Membranes were blocked 1h at RT under agitation in Intercept (TBS) Blocking Buffer (cat#927-60003, LI-COR Biosciences, Lincoln, NE, USA). Antibodies were diluted in 50% Intercept (TBS) Blocking Buffer + 45% DI-$H_2O$ + 5% TBS 10X (cat#1706435, Bio-Rad) + 0.1% Tween-20 (cat#P1379, MilliporeSigma). Membranes were incubated with primary antibodies overnight at 4°C in agitation, then washed 2x10min in TBS + 0.1% Tween-20. Membranes were incubated in secondary antibodies for 1h at RT in agitation and washed 4x10min in TBS + 0.1% Tween-20. Membranes were rinsed in TBS and imaged using an Odyssey CLx imaging system (LI-COR Biosciences). Antibodies and concentrations: 1:2000 anti-mCherry (cat#NBP2-25157, Novus Biological, Centennial, CO, USA), 1:3000 anti-α-tubulin, clone

DM1α (cat#T6199, Millipore Sigma), 1:5000 IRDye 680LT Donkey anti-Mouse IgG Secondary Antibody (cat#926-68022, LI-COR Biosciences), and 1:5000 IRDye 800CW Donkey anti-Rabbit IgG Secondary Antibody (cat#926-32213, LI-COR Biosciences). Raw data were quantified using FIJI with a consistent size Region-of-Interest within each image and protein of interest. The mean fluorescence intensity of background signal for each channel analysed was subtracted from the mean fluorescence intensity of the band of interest. For each sample, the mCherry mean fluorescence intensity was divided by the mean fluorescence intensity of the loading control (α-tubulin), then averaged across three replicates. Each replicate was run on separate gels and averaged for values seen in Supplementary Figure 10.

**Relative to Methods Section 4.8.** To evaluate the effects of compounds interfering with DOX induction in real-time, HeLa cells encoding DOX-inducible mCherry cells were plated at low density (2.5k/well) in a 96 well plate (cat#353072, Falcon by Corning) and imaged hourly with a 20X objective using the IncuCyte S3/SX1 G/R Optical Module (Red Acquisition Time=400msec) and the Adherent Cell-by-cell scan type. Cells were treated 5h with media +250ng/ml DOX, and washed with the same media in the presence of compound (3.33µM, 1.11µM, 370nM, 123nM, 41nM, or 14nM) or 0.33% DMSO (mock). As a readout, we measured red intensity (Total Integrated Intensity, RCU x µm2/image) within the objects identified by Adherent Cell-by-cell analysis (parameters used: Segmentation Adjustment=2, Cell Detection Sensitivity=1.6, Cell Contrast=3, Cell morphology=4; segmentation of the red channel was made with “Surface Fit No

Mask”). These data, together with the calibration described below, were used to normalize anastasis levels in compounds interfering with DOX induction.

A calibration was performed to correlate changes in anastasis with changes in mCherry (**Supplementary Figure 3**). HeLa cells with DOX-inducible caspase or DOX-inducible mCherry were plated at low density (3k/well) in a 96 well plate and imaged hourly with a 20X objective using the IncuCyte S3/SX1 G/R Optical Module (Green Acquisition Time=300msec; Red Acquisition Time=400msec) and the Adherent Cell-by-cell scan type. Cells were treated with DOX (75ng/ml, 150ng/ml, 250ng/ml, 500ng/ml or 1μg/ml) for 5h to transiently induce caspase expression, or continuously to induce mCherry expression. As a readout, we measured red intensity as described in above, and we calculated anastasis as described in **Methods Section 4.4.** For each timepoint imaged, we normalized anastasis or mCherry intensity by subtracting the lowest DOX concentration (75ng/ml). We then averaged across 3 replicates and smoothed the data out using a centered rolling average with a window of size 3. Next, for each timepoint, we trained a linear regression model using *polyfit* with a first degree polynomial from the *numpy* package, with the mCherry intensity differences as our input variable and anastasis differences as the response variables. After training the model for each timepoint (y = mx + b), we stored the trained coefficients (m, b).

To estimate how each compound influenced anastasis, we applied the model to the measured changes in red intensity for each treatment. The resulting predicted effects on anastasis were then incorporated into the kinome regularization screening curves by adding the predicted change to the original values. Original smoothed values were used for this adjustment when generating the final graphs, while raw original values were used for determining directionality, categorical classification, and regression output.

A comparable correction approach was applied to the induction dataset (see **Methods Section 4.7.2**), with two key modifications. First, calibration was based on the number of GC3AI$^+$ cells rather than anastasis values, and these counts were normalized to the lowest DOX concentration (75ng/ml). Second, the correction was applied to the induction curves themselves, rather than to the anastasis curves.

**Relative to Methods Section 4.9**. To correct treatment with fluorescence artifacts that interfered with our measurements, we developed an extrapolation-based strategy. For each compound, we began by measuring anastasis (**Methods Section 4.4**) and identifying the specific concentration(s) requiring extrapolation. For each timepoint across the full time course, we trained a simple linear regression model using the available concentrations that were not affected by the artifact. The predictor variable (*x*) was defined as the logarithm (base 3) of these concentrations, and the response (*y*) was the corresponding smoothed anastasis values. Once trained, this model was used to predict the expected anastasis level for the artifact-affected concentration(s), using the log-base-3 transformed concentration as input. The resulting predicted values then replaced the original measurements at those concentration points.

After extrapolation, the corrected dataset was processed differently depending on downstream use. For datasets prepared for kinome regularization analysis, all non-extrapolated measurements were reverted to raw, unsmoothed values, so that the extrapolated points were the only modeled elements. In contrast, when generating visualization plots, the smoothed dataset was retained. Standard deviation values were left unchanged.

**Relative to Methods Section 5.** Cells were imaged hourly with a 20X objective using the IncuCyte S3/SX1 G/R Optical Module (Green Acquisition Time=300msec) and the Adherent Cell-by-cell scan type. Masks to identify GC3AI$^{+}$ dead and living cells were made as described in **Methods Section 4.3**, and data were handled as described in **Methods Section 4.4**. Plots were generated using a custom python code and show the mean values and corresponding standard deviations processed using a centered three-point moving average (**Supplementary Figures 4-5**). To verify the effect of inhibitors, we reviewed images ~15h post-DOX or at the end of the experiment (~1d 5h post-DOX).

To calculate the % of GC3AI$^{+}$ cells (**Figure 6I-K**, **Supplementary Figure 4C**, **Supplementary Figure 8E**), we we quantified the total number of cells using the IncuCyte built-in Adherent Cell-by-cell analysis module (Cell boundary segmentation adjustment=2, Cell detection sensitivity=1.6, Cell contrast=3, Cell morphology=4). Importantly, high concentration of BSA (≥5%) and well coating with fibronectin reduced cell contrast and perturbed cells detection in samples that were not exposed to DOX, leading to an underestimation of the total number of cells, which could potentially result in an overestimation in the % of GC3AI$^{+}$ cells.

### Supplementary Figures

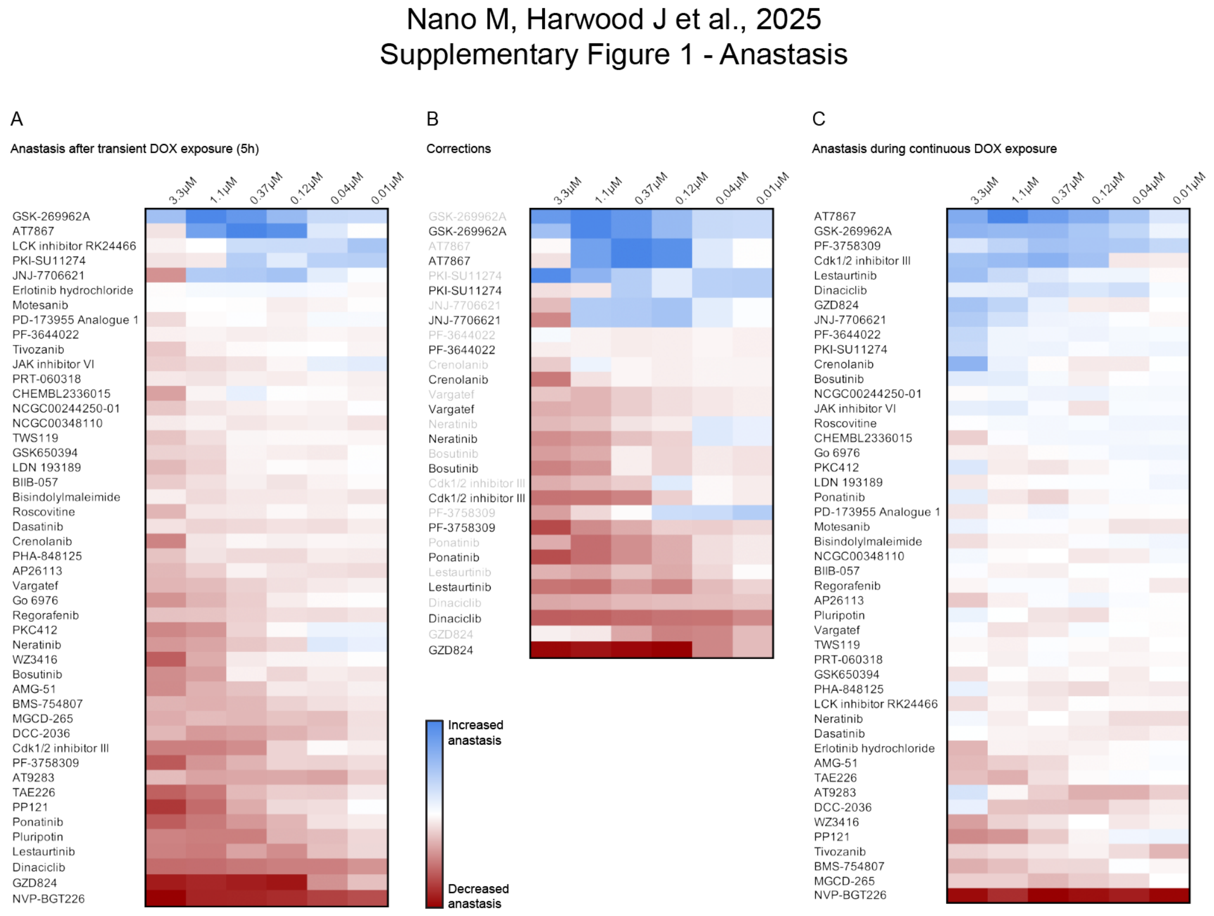


**Supplementary Figure 1. Kinase inhibitors modify anastasis.** Heatmaps showing anastasis following transient (A, B) or continuous (C) caspase activation. Anastasis was measured as the fraction of GC3AI⁺ cells surviving caspase activation, normalized to the mock control by subtraction, and averaged across unsmoothed values from time points 12–18 (n=2 biological replicates). **A.** Anastasis after corrections for artifacts and interference with DOX induction. **B.** Comparison of anastasis measures before (gray) and after (black) correction. **C.** Treatments increasing anastasis during continuous DOX exposure are likely to interfere with DOX induction. Increased anastasis is shown in blue, decreased anastasis is shown in red. Treatments that surpassed the threshold are shown in Supplementary Table 4.

Supplementary Figure 2 - Inhibitors effect without DOX

A

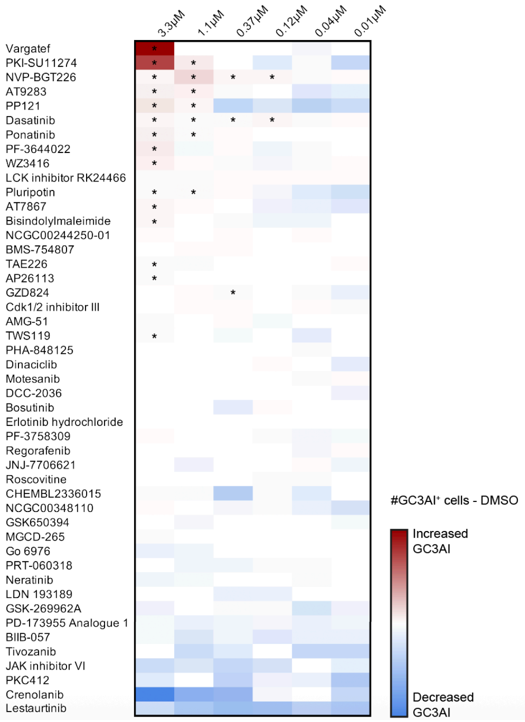


**Supplementary Figure 2. Inhibitors effect without DOX.** Heatmaps showing inhibitor effects in cells that were not treated with DOX. The number of GC3AI⁺ cells normalized to the mock control by subtraction, and averaged across unsmoothed values from time points 12–18 (n=2 biological replicates) is shown. Increased GC3AI is shown in red, decreased GC3AI is shown in blue. Asterisks label treatments increasing GC3AI above the toxicity threshold at least twice within the time window (Methods Section 4.6, Supplementary Table 3). No corrections were made (3.33 µM Vargatef, 3.33 µM PF-3644022, 3.33 µM and 1.11 µM PKI-SU11274 present green artifacts affecting the heatmap). JNJ-7706621, CHEMBL2336015, JAK3 inhibitor VI, NCG00244250-01, PD-173955 Analogue 1, Tivozanib, Roscovitine, PKC412, AMG-51, Go 6976, PRT-060318, Motesanib, BIIB-057, LDN 193189, Neratinib, Crenolanib, Bosutinib, DCC-2036, NCG00348110, Regorafenib, Cdk1/2 inhibitor III, PF-3758309, BMS-754807, PHA-848125, Dinaciclib, MGCD-265, and Lestaurtinib were only toxic in combination with with transient caspase activation (Table 2).

Nano M, Harwood J et al., 2025
Supplementary Figure 3 - Strategy to normalize interference with DOX induction

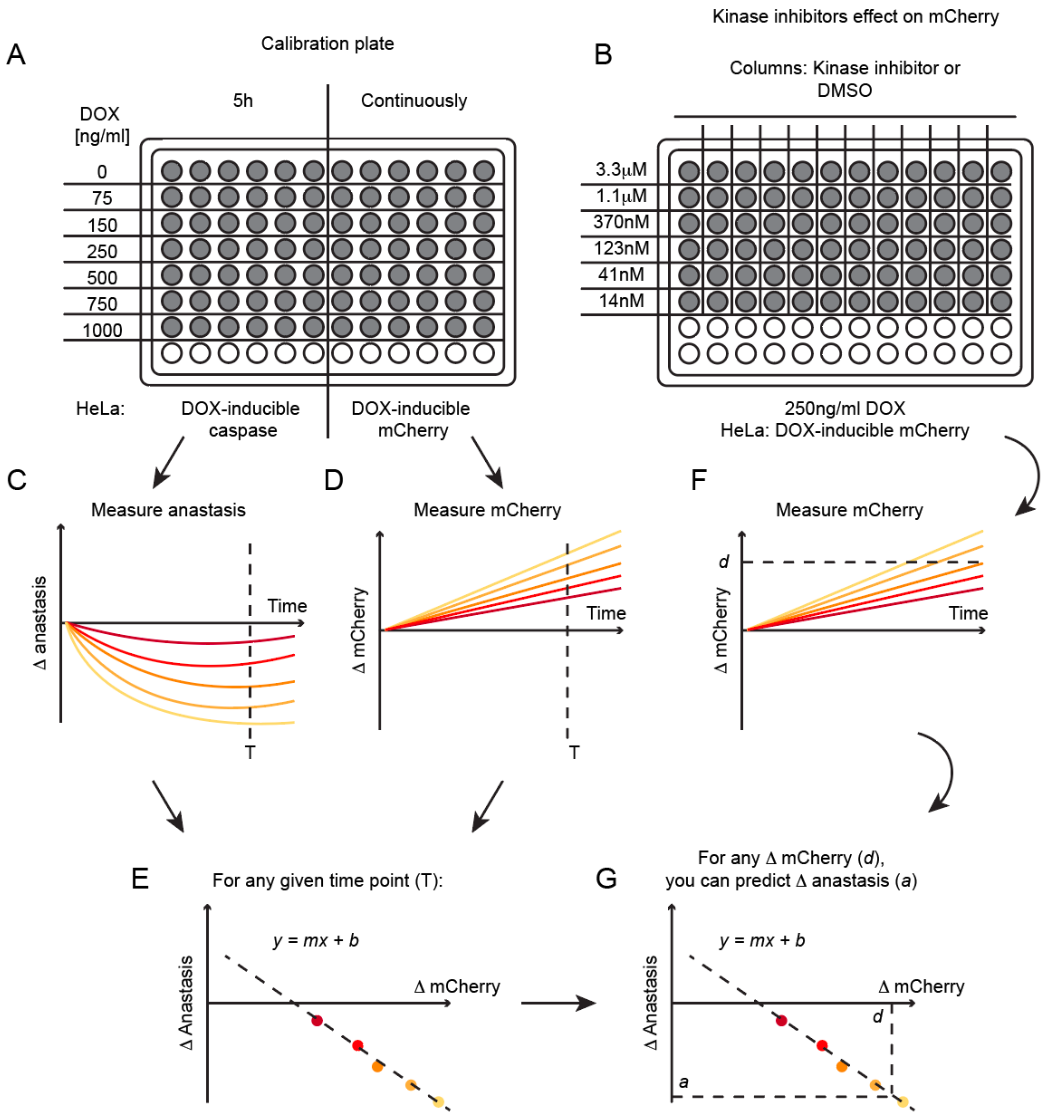


**Supplementary Figure 3. Strategy to normalize anastasis levels in compounds interfering with DOX induction.** We established a calibration between anastasis in cells encoding a DOX-inducible caspase and mCherry intensity in cells encoding a DOX-inducible mCherry by exposing both cell lines to a range of DOX concentrations (**A**). This calibration was then used to correct anastasis measurements in compounds that affected DOX induction in the screening dataset (Supplementary Figure 4). **A.** Cells expressing DOX-inducible caspase or DOX-inducible mCherry were treated with 0, 75, 150, 250, 500, 750, or 1000ng/ml DOX for 5 h (caspase) or continuously (mCherry). For each DOX concentration, we calculated and plotted the change in anastasis (Δ anastasis, **C**) and mCherry intensity (Δ mCherry, **D**) relative to 75 ng/ml DOX. We then plotted Δ anastasis versus Δ mCherry at each time point (T) and fit a linear regression (*y = m x + b*) (E). **B.**

To evaluate the effects of kinase inhibitors on DOX-inducible gene expression, cells expressing DOX-inducible mCherry were treated with 250 ng/ml DOX for 5 h, followed by 250 ng/ml DOX ± inhibitor at 3.3 µM, 1.1 µM, 370 nM, 123 nM, 41 nM, or 13 nM, or 0.33% DMSO. For each concentration, we quantified Δ mCherry (**F**), which was then used with the calibration in (**E**) to infer the expected Δ anastasis and normalize the screening results. **C-G** Schematic representations of hypothetical results and linear regression.

Nano M, Harwood J et al., 2025
Supplementary Figure 4 - Candidates screening

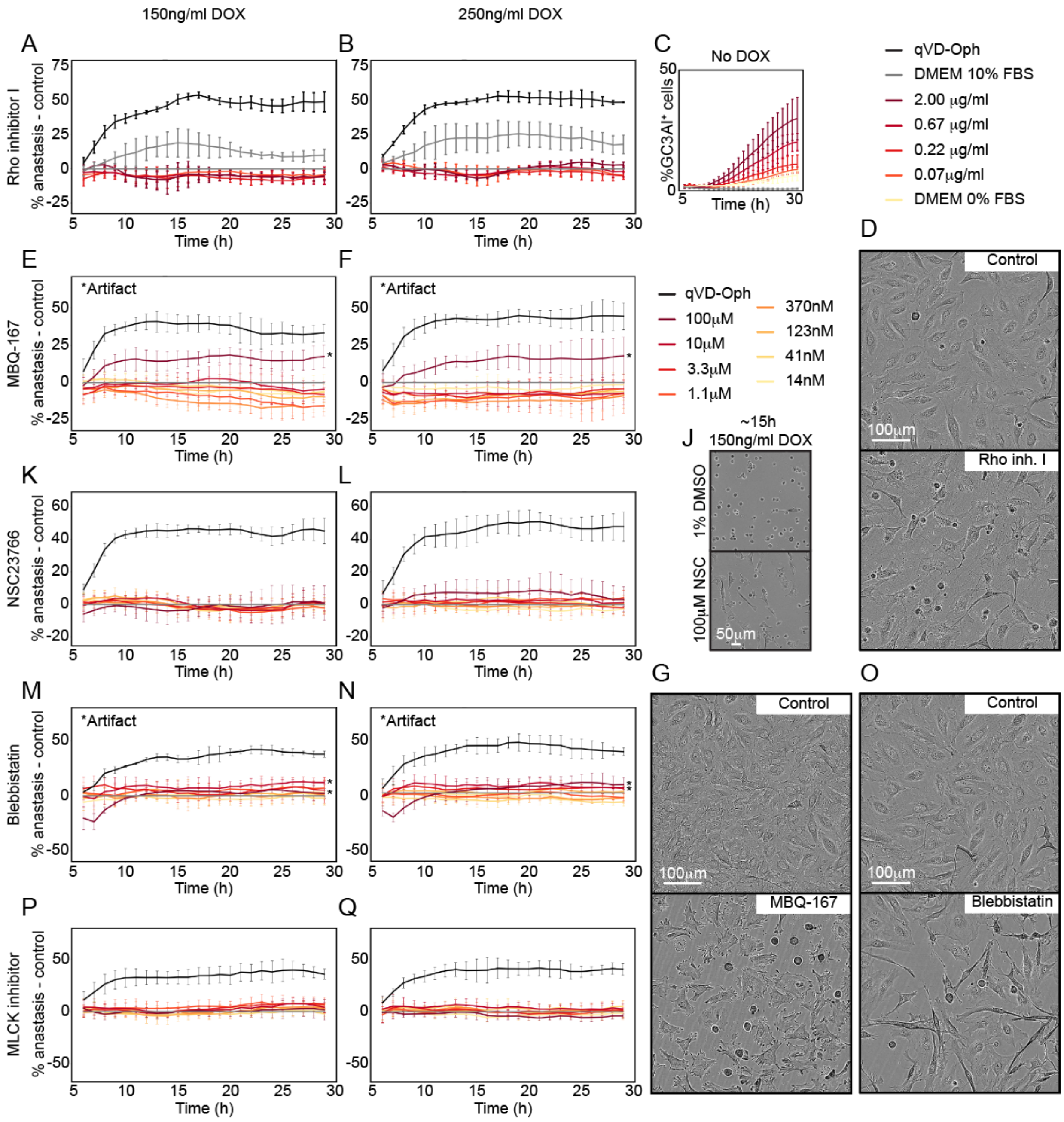


**Supplementary Figure 4. Anastasis in candidate compounds (Part I).** Plots and images illustrating the effects of different treatments. Plots show the percentage of anastasis following transient caspase activation normalized by subtraction to the control (A-B, E-F, K-L, M-N, P-Q), or the % of GC3AI⁺ cells (C). Data was normalized by subtraction and smoothed. The positive control (qVD-OPh) is shown in black. Different treatments are shown in distinct colors. Error bars represent the smoothed mean ± SD from at least 2 independent experiments. **A-B.** Rho inhibitor I did not affect anastasis in cells with medium-high levels of caspase activation. **C.** In samples that were not exposed to DOX, Rho inhibitor I increased the fraction of GC3AI$^+$ cells. **D.** Cells grown in basal media (control) or basal media + 2µg/ml Rho inhibitor I at the end of our experiment. **E-G**. MBQ-167 caused a variable reduction in survival with caspase activation as well as morphological alterations and mitotic arrest (E-F). At 100µM, the compound precipitated out of solution, forming green crystals that perturbed the automated quantifications (*Artifact). On its own, MBQ-167 induced high rates of mitotic arrest (G). **G.** Cells grown in complete media 0.1% DMSO (control) or complete media + 10µM MBQ-167 at the end of our experiment. **J-L.** NSC23766 had little impact on anastasis (K-L), although at high concentrations it appeared to interfere with DOX induction, as indicated by the presence of living cells under continuous DOX exposure (J). **M-O.** Blebbistatin. At 100 µM, and to a lesser extent at 10 µM, blebbistatin was toxic both in the presence of caspase activation (M-N) and when applied alone (O); however, the compound precipitated out of solution, forming green crystals that interfered with automated quantification (*Artifact). At lower concentrations, the effects on survival were negligible. **O.** Cells grown in complete media 0.1% DMSO (control) or complete media + 10µM Blebbistatin at the end of our experiment. **P-Q.** MLCK inhibitor peptide 18.

Nano M, Harwood J et al., 2025
Supplementary Figure 5 - Candidates screening

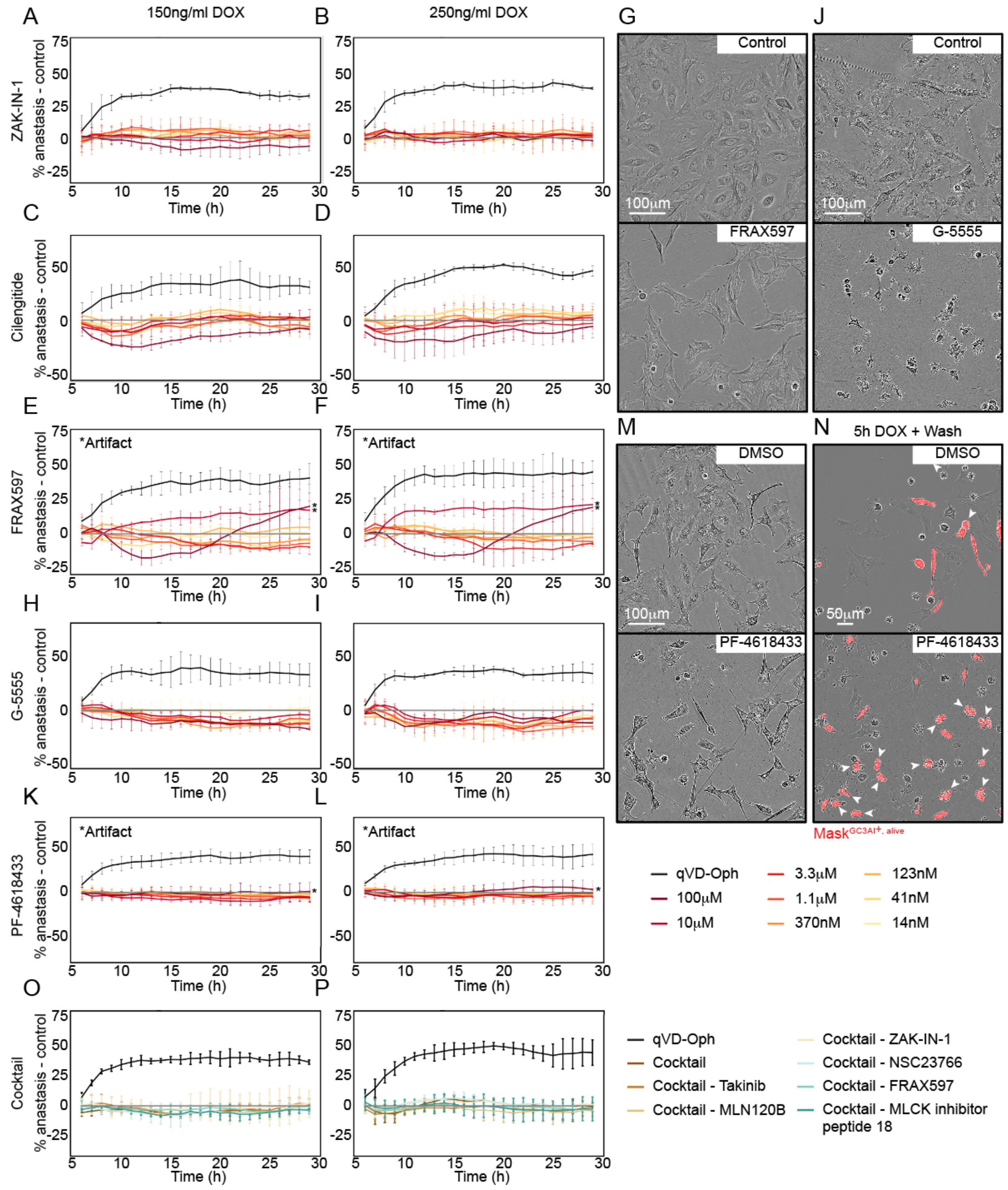


**Supplementary Figure 5. Anastasis in candidate compounds (Part II) and cocktail of candidate compounds.** Plots and images illustrating the effects of different treatments. Plots show the percentage of anastasis following transient caspase activation normalized to the control for cells treated with 150ng/ml (A, C, E, H, K, O) or 250ng/ml (B, D, F, I, L, P) DOX. Data were normalized by subtraction and smoothed. The positive control (qVD-OPh) is shown in black. Different treatments are shown in distinct colors. Error bars represent the smoothed mean ± SD from at least 2 independent experiments. **A-B.** ZAK-IN-1. **C-D.** Cilengitide considerably reduces survival at 100µM, both in the presence (C-D) and absence (not shown) of caspase activation. **E-G.** FRAX597 caused a small, variable reduction in survival with caspase activation (E-F). It also reduced survival on its own at higher concentrations (G). At 100µM and 10µM, the compound precipitated out of solution, forming green crystals that perturbed automated quantifications (*Artifact). **G.** Cells grown in complete media 0.1% DMSO (control) or complete media + 3.3µM FRAX597 at the end of our experiment. **H-J.** At concentrations ≤10µM, G-5555 caused a modest reduction in survival after transient caspase activation (H-I). Abnormal dying cell morphology, accompanied by loss of GC3AI fluorescence led to a gross underestimation of cell death caused by 100µM G-5555 (~100%). **J.** Cells grown in complete media 0.1% DMSO (control) or complete media 100µM G-5555 at the end of our experiment. On its own, G-5555 only caused cell death when applied at 100µM. **K-N.** PF-4618433 has a negligible effect on cell survival at all concentrations below 100µM. The abnormal morphology of dying cells causes the quantifications to underestimate the frequency of cell death caused by 100µM PF-4618433 (*Artifact). **M.** Cells grown in complete media 1% DMSO (control) or complete media + 100µM PF-4618433 at the end of our experiment. Despite a potential interference with DOX induction, 100µM PF-4618433 caused mitotic arrest, and inhibited survival both in the presence of caspase and on its own. **N.** Cells were transiently exposed to DOX (5h, 150 ng/ml) and then washed with either complete medium containing 1% DMSO (control) or with 100 µM PF-4618433. Images are relative to 15h after DOX treatment. The mask used to identify GC3AI-positive, living cells is shown in red. Treatment with 100 µM PF-4618433 caused dying cells to adopt an abnormal morphology, leading to misclassification (white arrowheads). **O-P.** Plots showing the percentage of anastasis following transient caspase activation, normalized to the DMSO control, for cells treated with qVD-OPh, a cocktail of candidate compounds (Takinib, MLN120B, ZAK-IN-1, NSC23766, FRAX597, and MLCK inhibitor peptide 18; 250 nM each), or the same cocktail lacking one compound at a time.

Supplementary Figure 6 - Growth factor signaling

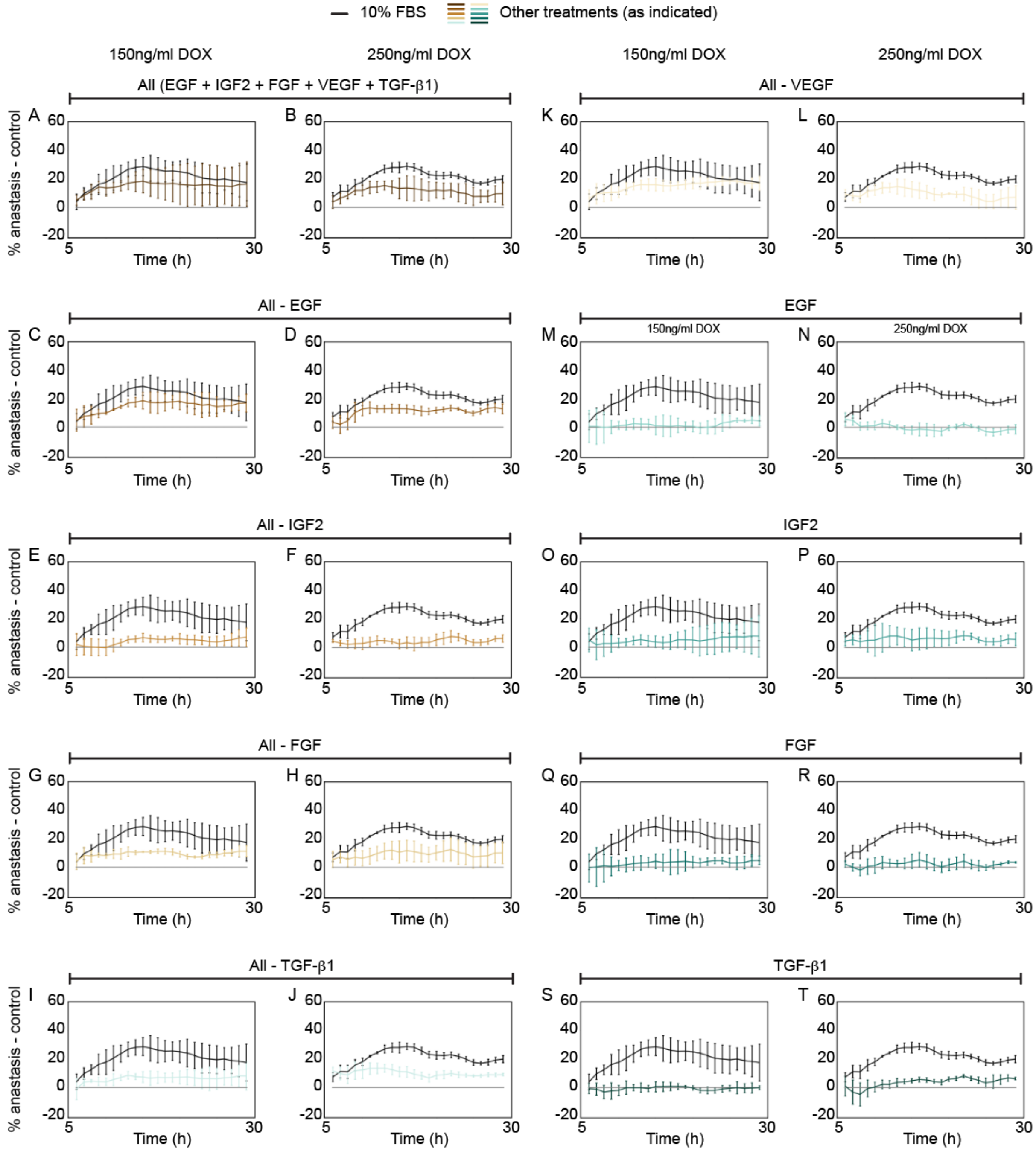


**Supplementary Figure 6. Growth factor signaling supports recovery from direct effector caspase activation (Part I).** % anastasis normalized over the control (base media, FBS-free, gray) for cells washed with complete media (black curve) or base media with different growth factors combinations. This figure shows the same data reported in Figure 5, separated according to the combination of growth factors used in the wash for cells transiently exposed to 150ng/ml DOX

(A, C, E, G, I, K, M, O, Q, S), or 250ng/ml DOX (B, D, F, H, J, L, N, P, R, T). The effect of IGF2, though modest in this experiment, was larger in others (see **Supplementary Figure 7**). In each graph, the black curve represents media containing 10% FBS. Data and error bars were smoothed. Error bars represent the mean ± SD from n=2 independent experiments.

Nano M, Harwood J et al., 2025
Supplementary Figure 7 - Growth factor signaling

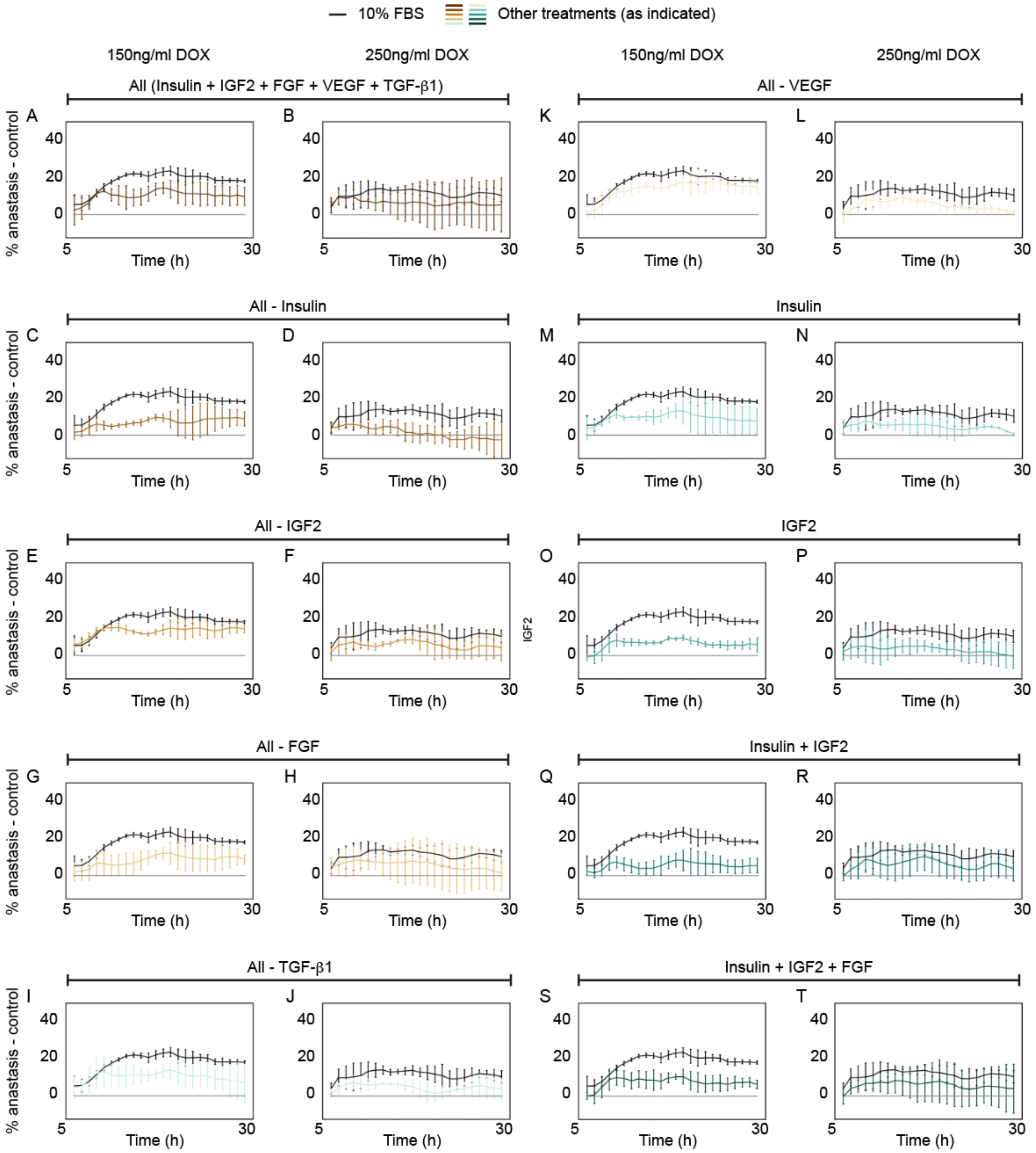


**Supplementary Figure 7. Growth factor signaling supports recovery from direct effector caspase activation (Part II).** % anastasis normalized over the control (base media, FBS-free, gray) for cells washed with complete media (black curve) or base media with different growth factors combinations. This figure shows the same data reported in Figure 5, separated according to the combination of growth factors used in the wash for cells transiently exposed to 150ng/ml DOX (A, C, E, G, I, K, M, O, Q, S), or 250ng/ml DOX (B, D, F, H, J, L, N, P, R, T). In each graph, the black curve represents media containing 10% FBS. The effect size for IGF2 varied between experiments (Supplementary Figure 6). Data and error bars were smoothed. Error bars represent the mean ± SD from n=2 independent experiments.

Nano M, Harwood J et al., 2025
Supplementary Figure 8 - Fibronectin, collagen, and matrigel coating

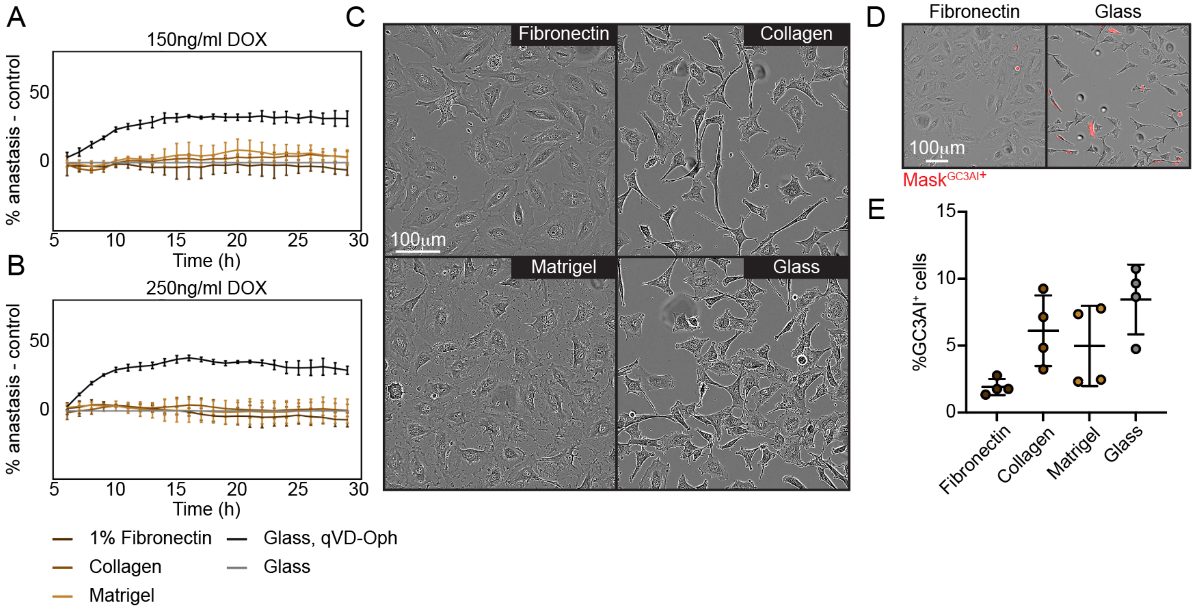


**Supplementary Figure 8. Neither fibronectin, collagen, nor matrigel affected iCasp3-GC3AI anastasis. A-B.** Plots showing the percentage of anastasis normalized to the control (glass) for cells plated on different substrates and transiently exposed to DOX (5h). Data were normalized by subtraction and smoothed. Error bars represent the mean ± SD (smoothed) from n=2 independent experiments. **C-D.** Phase contrast images acquired at the Incucyte for cells plated on different substrates before DOX-treatment. **D.** Cells recognized as GC3AI$^+$ are labeled in red (see Methods Section 5). **E.** Dot plot showing the % of GC3AI$^+$ cells on different substrates before DOX-treatment. n=4 independent experiments. Error bars=mean ± SD.

Supplementary Figure 9 - GC3AI induction in samples transiently exposed to DOX

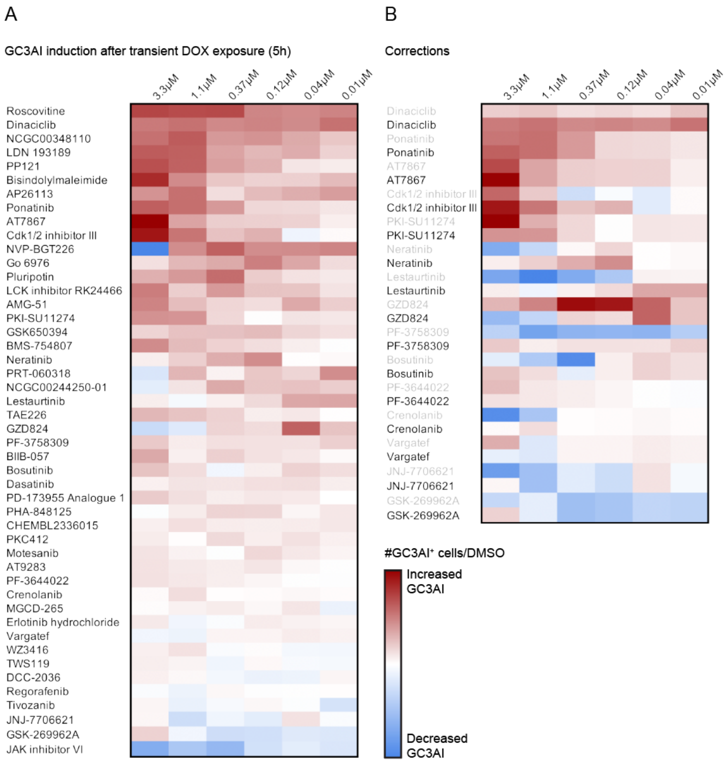


**Supplementary Figure 9. GC3AI induction in samples transiently exposed to DOX.** Heatmaps showing GC3AI induction following 5h of DOX exposure calculated as the ratio of GC3AI$^+$ cells to DMSO and averaged across unsmoothed values from time points 12-18 (n=2 biological replicates). **A.** Induction corrected for artifacts and interference with DOX induction. **B.** Comparison of induction before (gray) and after (black) correction. Increased induction is shown in red, decreased induction is shown in blue.

Supplementary Figure 10 - Western Blot to assess the effect of inhibitors on DOX induction

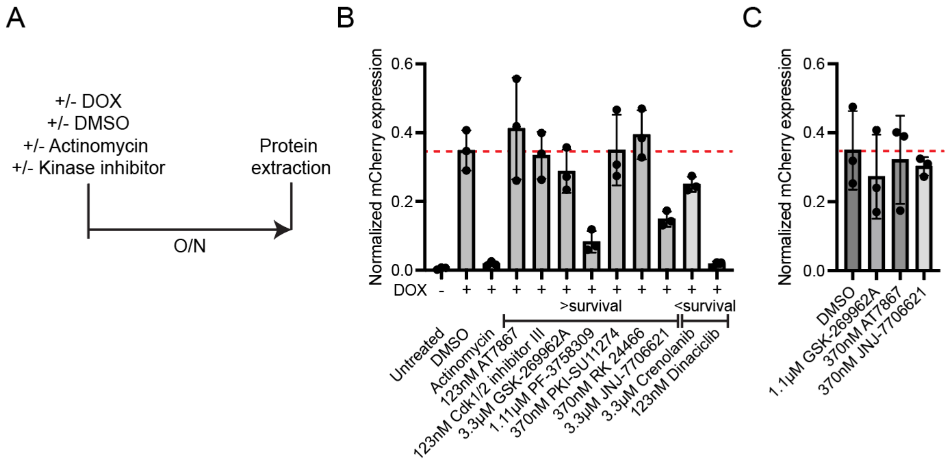


**Supplementary Figure 10. Western Blot to assess the effect of inhibitors on DOX induction.** HeLa DOX-mCherry cells were treated with DOX and kinase inhibitors to establish any interference with DOX induction. **A.** HeLa DOX-mCherry cells were treated overnight (O/N) with 1 µg/ml DOX and actinomycin D, DMSO, or one concentration of kinase inhibitor before lysis the following morning (Methods 4.7.3). **B-C.** Bar graphs showing mCherry protein intensity normalized to α-tubulin for the indicated treatments. The red dotted line indicates baseline mCherry expression in the DMSO control. Error bars represent mean ± SD from n=3 independent experiments.

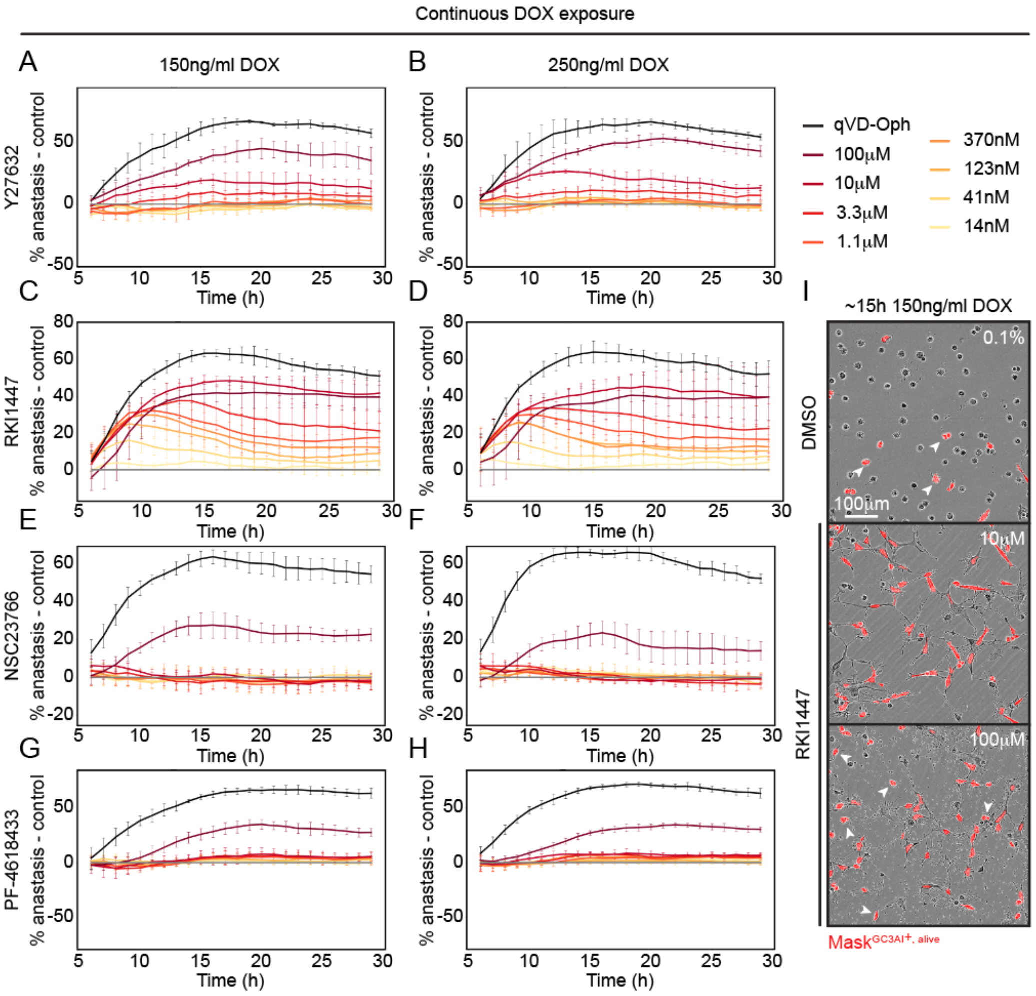


**Supplementary Figure 11. Candidate screening: inhibitors interfering with DOX induction. A-H.** Plots showing anastasis for inhibitors that appear to interfere with DOX induction, as inferred from the presence of surviving cells during continuous DOX exposure. **I.** High concentrations of inhibitors can lead to morphological alteration perturbing the detection of live cells. Cells exposed to 150ng/DOX for ~15h that were treated with 0.1% DMSO or RKI1447 at the 5h mark. The mask used to identify GC3AI-positive, living cells is shown in red. Treatment with high concentrations of RKI1447 caused dying cells to adopt an abnormal morphology, leading to misclassification. White arrowheads point to examples of misclassification. Similar morphological alterations were observed for cells exposed to Y27632 (not shown).

### Supplementary Tables

### Supplementary Table 1.

| CAS | Compound name and notes | ID |
|---|---|---|
| 890019-63-3 | AMG-51 | NCGC00263201 |
| 1197958-12-5 | AP26113 | NCGC00351602 |
| 857531-00-1 | AT7867 | NCGC00346549 |
| 896466-04-9 | AT9283 | NCGC00346493 |
| 1370261-96-3 | BIIB-057 | NCGC00351606 |
| 133052-90-1 | Bisindolylmaleimide | NCGC00262009 |
| 1001350-96-4 | BMS-754807 | NCGC00346453 |
| 380843-75-4 | Bosutinib | NCGC00241107 |
| 443798-55-8 | Cdk1/2 Inhibitor III | NCGC00387780 |
| 1429617-90-2 | CHEMBL2336015 | NCGC00347217 |
| 670220-88-9 | Crenolanib | NCGC00346658 |
| 302962-49-8 | Dasatinib | NCGC00181129 |
| 1020172-07-9 | DCC-2036 | NCGC00263172 |
| 779353-01-4 | Dinaciclib | NCGC00346656 |
| 183319-69-9 | Erlotinib hydrochloride | NCGC00164574 |
| 136194-77-9 | Go 6976 | NCGC00163451 |
| 850664-21-0 | GSK-269962A | NCGC00241982 |
| 890842-28-1 | GSK650394 | NCGC00250410 |
| 1421783-64-3 | GZD824 | NCGC00351607 |
| 856436-16-3 | JAK3 Inhibitor VI | N/A |
| 443797-96-4 | JNJ-7706621 | NCGC00263151 |
| 1062368-24-4 | LDN 193189 | NCGC00249389 |
| 111358-88-4 | Lestaurtinib | NCGC00244256 |
| 875337-44-3 | MGCD-265 | NCGC00483925 |
| 453562-69-1 | Motesanib | NCGC00263205 |
| 1261496-45-0 | NCGC00244250-01 | NCGC00244250 |
| NCGC00348110 | NCGC00348110 | NCGC00348110 |
| 698387-09-6 | Neratinib | NCGC00241101 |
| 1245537-68-1 | NVP-BGT226 | NCGC00346681 |

| | | |
|---|---|---|
| 185039-990 | PD-173955 Analogue 1 | NCGC00345830 |
| 1276121-88-0 | PF-3644022 | NCGC00351595 |
| 898044-15-0 | PF-3758309 | NCGC00345784 |
| 802539-81-7 | PHA-848125 | NCGC00346673 |
| 120685-11-2 | PKC 412 | NCGC00241102 |
| 658084-23-2 | PKI-SU11274 | NCGC00165902 |
| 839707-37-8 | Pluripotin | NCGC00167403 |
| 943319-70-8 | Ponatinib | NCGC00263152 |
| 1092788-83-4 | PP121 | NCGC00346619 |
| 1194961-19-7 | PRT-060318 | NCGC00253909 |
| 755037-03-7 | Regorafenib | NCGC00263138 |
| 213743-31-8 | RK 24466 | NCGC00015280 |
| 186692-46-6 | Roscovitine (Seliciclib) | NCGC00094374 |
| 761437-28-9 | TAE226 | NCGC00346931 |
| 475108-18-0 | Tivozanib | NCGC00249390 |
| 601514-19-6 | TWS119 | NCGC00346638 |
| 928326-83-4 | Vargatef | NCGC00263156 |
| 1214265-56-1 | WZ3416 | NCGC00346496 |

**Supplementary Table 1.** List of inhibitors used in our screening. The drugs were selected to inhibit (result in <0.5 residual activity) at least 10% of the human kinome at the highest concentration (3.3µM). JAK3 Inhibitor VI and MGCD-265 were screened but excluded from the kinome regularization. Cdk1/2 Inhibitor III: we screened compound CAS#443798-47-8 instead of CAS#443798-55-8. LDN 193189: we were unable to fully dissolve the compound.

**Supplementary Table 2.**

| Compound | 3.3μM | 1.1μM | 0.37μM | 0.12μM | 0.04μM | 0.01μM |
|---|---|---|---|---|---|---|
| Cdk1/2 inhibitor III | xx | xx | xx | xx | xx | xx |
| LCK inhibitor RK24466 | - | - | - | - | - | ✓✓ |
| PKC412 | xx | xx | xx | - | ✓✓ | - |
| JAK inhibitor VI | xx | xx | - | - | - | ✓✓ |
| Vargatef | xx | xx | xx | xx | xx | xx |
| NCGC00244250-01 | xx | xx | - | - | - | - |
| AMG-51 | xx | xx | xx | - | - | - |
| Go 6976 | xx | xx | xx | - | - | - |
| PRT-060318 | xx | xx | - | - | - | xx |
| Bosutinib | xx | xx | - | xx | xx | xx |
| NVP-BGT226 | xx | xx | xx | xx | xx | xx |
| JNJ-7706621 | xx | - | ✓✓ | ✓✓ | ✓✓ | - |
| WZ3416 | xx | xx | xx | - | xx | xx |
| GSK650394 | - | - | - | - | - | - |
| AT7867 | xx | ✓✓ | ✓✓ | ✓✓ | - | - |
| Motesanib | - | - | x | xx | x | - |
| PD-173955 Analogue 1 | | x | - | - | - | - |
| PP121 | xx | xx | xx | xx | xx | ✓ |
| Pluripotin | xx | xx | xx | xx | xx | xx |
| PF-3758309 | xx | xx | xx | xx | xx | xx |
| Neratinib | xx | xx | xx | xx | - | - |
| Tivozanib | xx | xx | - | - | - | - |
| Dasatinib | - | - | - | - | xx | xx |
| Crenolanib | xx | xx | - | xx | - | |
| TWS119 | xx | xx | xx | - | - | - |
| AT9283 | xx | xx | xx | xx | xx | xx |
| BMS-754807 | xx | xx | xx | xx | xx | xx |
| Erlotinib hydrochloride | - | - | - | - | - | - |
| PHA-848125 | xx | xx | xx | xx | xx | xx |
| Dinaciclib | xx | xx | xx | xx | xx | xx |
| DCC-2036 | xx | xx | xx | xx | xx | - |
| Roscovitine | xx | xx | - | - | - | - |
| Bisindolylmaleimide | - | xx | xx | xx | xx | xx |
| NCGC00348110 | xx | - | xx | xx | xx | xx |
| BIIB-057 | xx | xx | - | - | x | - |
| AP26113 | xx | xx | xx | - | - | - |

| | | | | | | |
|---|---|---|---|---|---|---|
| MGCD-265 | xx | xx | xx | xx | xx | xx |
| GSK-269962A | ✓✓ | ✓✓ | ✓✓ | ✓✓ | - | - |
| GZD824* | xx | xx | xx | xx | xx | xx |
| Ponatinib | xx | xx | xx | xx | xx | xx |
| LDN 193189 | xx | xx | x | - | - | - |
| CHEMBL2336015 | | - | - | - | - | - |
| TAE226 | xx | xx | xx | xx | xx | - |
| PF-3644022 | - | - | - | - | - | xx |
| PKI-SU11274 | xx | xx | ✓✓ | - | ✓✓ | ✓✓ |
| Regorafenib | xx | xx | xx | - | xx | xx |
| Lestaurtinib | xx | xx | xx | xx | xx | xx |

**Supplementary Table 2.** Treatments' effect on anastasis within the 12-18 hour window. Related to Table 1 and Table 2. (xx) indicates -1, (x) -0.5, (-) 0, (✓) +0.5, (✓✓) +1. *3.33 and 1.11 µM GZD824 were corrected by manual counting (see Methods Section 4.9.1).

**Supplementary Table 3**.

| | 3.3µM | 1.1µM | 0.37µM | 0.12µM | 0.04µM | 0.01µM |
|---|---|---|---|---|---|---|
| Dasatinib | x | x | x | x | | |
| NVP-BGT226 | x | x | x | x | | |
| AT9283 | x | x | | | | |
| Pluripotin | x | x | | | | |
| PKI-SU11274* | x | x | | | | |
| Ponatinib | x | x | | | | |
| PP121 | x | x | | | | |
| AP26113 | x | | | | | |
| AT7867 | x | | | | | |
| Bisindolylmaleimide | x | | | | | |
| GZD824 | | | x | | | |
| PF-3644022 | x | | | | | |
| TAE226 | x | | | | | |
| TWS119 | x | | | | | |
| Vargatef* | x | | | | | |
| WZ3416 | x | | | | | |

**Supplementary Table 3**. Summary of treatments that increased GC3AI activation in the absence of caspase activation within the 12-18h time window after data normalization and smoothing. Treatments were considered positive only if GC3AI signal exceeded the threshold at ≥2 time points within this window and are indicated in red and marked with a [illegible]. *Increased GC3AI activation due to compound autofluorescence.

**Supplementary Table 4.**

| | 3.3µM | 1.1µM | 0.37µM | 0.12µM | 0.04µM | 0.01µM |
|---|---|---|---|---|---|---|
| AT7867 | ✓ | ✓ | ✓ | ✓ | ✓ | x |
| Dinaciclib | ✓ | ✓ | ✓ | ✓ | ✓ | x |
| GSK-269962A | ✓ | ✓ | ✓ | ✓ | ✓ | x |
| Cdk1/2 inhibitor III | ✓ | ✓ | ✓ | ✓ | x | x |
| PF-3758309 | x | ✓ | ✓ | ✓ | ✓ | x |
| JNJ-7706621 | ✓ | ✓ | x | x | x | x |
| Crenolanib | ✓ | x | x | x | x | x |
| GZD824 | ✓ | x | x | x | x | x |
| Lestaurtinib | ✓ | x | x | x | x | x |
| PF-3644022 | ✓ | x | x | x | x | x |
| PKI-SU11274 | ✓ | x | x | x | x | x |

**Supplementary Table 4.** List of treatments increasing the fraction of living cells in samples continuously exposed to DOX. Note that some of these compounds have toxic properties, and do not improve survival in cells transiently exposed to DOX.

**Supplementary Table 5.**

| | 3.3μM | 1.1μM | 0.37μM | 0.12μM | 0.04μM | 0.01μM |
|---|---|---|---|---|---|---|
| CHEMBL2336015 | xx | xx | xx | xx | xx | xx |
| Dinaciclib | xx | xx | xx | xx | xx | xx |
| LCK inhibitor RK 24466 | xx | xx | xx | xx | xx | xx |
| NCGC00348110 | xx | xx | xx | xx | xx | xx |
| PP121 | xx | xx | xx | xx | xx | xx |
| NCGC00244250-01 | - | xx | xx | xx | xx | xx |
| PHA-848125 | - | xx | xx | xx | xx | xx |
| Pluripotin | xx | xx | xx | xx | - | xx |
| Ponatinib | xx | xx | xx | xx | xx | - |
| TAE226 | xx | xx | xx | xx | xx | - |
| AT7867 | xx | xx | xx | xx | - | - |
| BMS-754807 | xx | xx | xx | xx | - | - |
| Cdk1/2 inhibitor III | xx | xx | xx | xx | - | - |
| Dasatinib | - | xx | xx | xx | xx | - |
| Lestaurtinib | xx | - | - | xx | xx | xx |
| Motesanib | xx | - | - | xx | xx | xx |
| PD-173955 Analogue 1 | xx | xx | xx | - | xx | - |
| PKC412 | xx | - | xx | - | xx | xx |
| Roscovitine | xx | xx | - | - | xx | xx |
| AMG-51 | xx | - | xx | - | xx | - |
| Bisindolylmaleimide | xx | xx | - | - | - | xx |
| LDN 193189 | xx | xx | xx | - | - | - |
| Neratinib | - | xx | xx | xx | - | - |
| PRT-060318 | - | xx | - | xx | - | xx |
| AP26113 | xx | xx | - | - | - | - |
| Go 6976 | - | - | xx | - | xx | - |
| GSK650394 | - | - | - | xx | xx | - |
| PKI-SU11274 | xx | xx | - | - | - | - |
| Crenolanib | x | xx | - | - | - | - |
| Bosutinib | xx | - | - | - | - | - |

| | | | | | | |
|---|---|---|---|---|---|---|
| AT9283 | - | - | - | - | - | - |
| BIIB-057 | - | - | - | - | - | - |
| PF-3644022 | - | - | - | - | - | - |
| PF-3758309 | - | - | - | - | - | - |
| Regorafenib | - | - | - | - | - | - |
| Vargatef | - | - | - | - | - | - |
| NVP-BGT226 | ✓✓ | - | xx | xx | xx | xx |
| MGCD-265 | - | - | - | xx | xx | ✓✓ |
| DCC-2036 | xx | - | - | - | ✓✓ | - |
| Erlotinib hydrochloride | xx | ✓✓ | - | - | - | - |
| GZD824 | xx | xx | ✓✓ | ✓✓ | xx | xx |
| WZ3416 | xx | xx | ✓✓ | - | ✓✓ | ✓✓ |
| JNJ-7706621 | xx | ✓✓ | ✓✓ | ✓✓ | - | - |
| Tivozanib | xx | ✓✓ | - | - | - | ✓✓ |
| JAK3 inhibitor VI | ✓✓ | ✓✓ | ✓✓ | ✓✓ | ✓✓ | ✓✓ |
| GSK-269962A | - | - | ✓✓ | ✓✓ | - | ✓✓ |
| TWS119 | - | - | ✓✓ | - | - | - |

**Supplementary Table 5.** Summary of GC3AI induction in samples transiently exposed to DOX relative to the 12-18h time window after data normalization and correction. Treatments increasing the fraction of GC3AI$^+$ cells are shown in red. Treatments decreasing the fraction of GC3AI$^+$ cells are shown in blue. Neutral treatments are shown in white. (xx) indicates -1, (x) -0.5, (-) 0, (✓) +0.5, (✓✓) +1. Relative to **Supplementary Figure 9**.

**Supplementary Table 6. Rationale used to determine treatments to test by western blot.**

| Inhibitor | Concentration | Rationale |
|---|---|---|
| Cdk1/2 Inhibitor III | 123nM | Only concentration potentially enhancing anastasis |
| PKI-SU11274 | 370nM | Highest, artifact-free concentration potentially enhancing anastasis |
| RK 24466 | 370nM | Highest concentration potentially enhancing anastasis |
| AT7867 | 370nM | Intermediate concentration promoting survival in unwashed samples |
| Crenolanib | 3.3µM | Highest concentration promoting survival in unwashed samples |
| GSK-269962A | 3.3µM | Highest concentration promoting survival in unwashed samples |
| JNJ-7706621 | 3.3µM | Highest concentration promoting survival in unwashed samples |
| PF-3758309 | 1.1µM | Medium-high concentration promoting survival in unwashed samples |
| Dinaciclib | 123nM | Medium-low concentration promoting survival in unwashed samples |

**Supplementary Table 7. List of corrected treatments.**

| Compound | Concentrations (µM) |
|---|---|
| Cdk1/2 Inhibitor III | 3.3, 1.1, 0.37, 0.12 |
| Dinaciclib | 3.3, 1.1, 0.37, 0.12, 0.04, 0.01 |
| JNJ-7706621 | 3.3 |
| Lestaurtinib | 3.3, 1.1, 0.37, 0.12, 0.04, 0.01 |
| Neratinib | 3.3, 1.1, 0.37, 0.12 |
| PF-3758309 | 3.3, 1.1, 0.37, 0.12, 0.04, 0.01 |
| Crenolanib | 3.3, 1.1 |
| PKI-SU11274 | 3.3, 1.1 |
| Bosutinib | 3.3, 1.1, 0.37* |
| GSK-269962A | 3.3 |
| AT7867 | 3.3 |

*Only modified for induction, not anastasis

**Supplementary Table 8.**

| Inhibitor | Source | Target |
|---|---|---|
| Rho inhibitor I | cat#CT04-A, Cytoskeleton Inc., Denver, CO, USA | Rho (1) |
| MBQ-167 (CAS: 2097938-73-1) | cat#HY-112842, MedChemExpress, Monmouth Junction, NJ, USA | Rac/Cdc42 (2) |
| **NSC23766 trihydrochloride (CAS: 1177865-17-6) | cat#A1952, ApexBio | Rac (3) |
| **PF-4618433 (CAS: 1166393-85-6) | cat#HY-18312, MedChemExpress | FAK2/PTK2B (4) |
| Chroman I (CAS: 1273579-40-0) | cat#38248, Cayman Chemicals | ROCK (5) |
| *Y-27632 dihydrochloride (CAS: 129830-38-2) | cat#HY-10583, MedChemExpress | ROCK (6) |
| *RKI-1447 (CAS: 1342278-01-6) | cat#HY-15755, MedChemExpress | ROCK (7) |
| Blebbistatin (CAS: 856925-71-8) | cat#HY-13441, MedChemExpress | Myosin (8) |
| MLCK Inhibitor Peptide 18 (CAS: 224579-74-2) | cat#19181, Cayman Chemicals | Myosin Light Chain Kinase (9) |
| ZAK-IN-1 (CAS: 2362525-64-0) | cat#HY-128326, MedChemExpress | Zak (10,11) |
| Cilengitide (CAS: 188968-51-6) | cat#77235-020, Avantor, Radnor, PA, USA | Integrins (12) |
| FRAX597 (CAS: 1286739-19-2) | cat#HY-15542A, MedChemExpress | Focal adhesion dynamics (PAK1/2/3) (13) |
| G-5555 (CAS: 2319590-15-1) | cat#HY-19635A, MedChemExpress | Focal adhesion dynamics (PAK1/2) (14) |

**Supplementary Table 8.** List of inhibitors tested in our candidate screening, their source, and their main targets. *Compounds were not evaluated because they interfered with DOX-induction, as determined by a reduction in cell death in samples continuously exposed to DOX, and/or altered cell morphology, preventing accurate detection of living cells (Supplementary Figure 11). **Compounds were evaluated despite their highest concentration interference with DOX-induction (Supplementary Figure 11).

**Supplementary Table 9.**

| Growth factor | Origin | Final concentration used | Physiological levels | Manufacturer ED50 recommendation |
|---|---|---|---|---|
| EGF | cat#236-EG-200, R&D systems | 100ng/ml | ~0.5-87ng/ml (15) | ED50=0.02-0.1 ng/ml in a Balb/3T3 proliferation assay (up to 50ng/ml for intestinal organoid culture) |
| IGF2 | cat#292-G2-050, R&D systems | 1000ng/ml | ~400-1000ng/ml (16,17) | ED50=1.5-6 ng/ml in a serum-free cell proliferation assay using MCF-7 |
| FGF | cat#BT-FGFBHS, R&D systems (17) | 500ng/ml | ~0.004-0.01 ng/ml (18) | ED50=0.05-0.6 ng/ml in a cell proliferation assay using NR6R-3T3 (up to 100 ng/ml in Human iPSCs culture) |
| VEGF | cat#BT-VEGF-020, R&D systems | 100ng/ml | ~0.08-0.5 ng/ml (18,19) | ED50=1.50-12.0 ng/ml in a cell proliferation assay using HUVEC |
| TGF-β1 | cat#7754-BH-005/CF, R&D systems | 100ng/ml | ~0.5-25ng/ml (20) | ED50=0.04-0.2 ng/ml measured by the ability to inhibit the IL-4-dependent proliferation of HT-2 mouse T cells |
| Insulin | cat#I9278, Millipore Sigma | 1000μg/mL | ~$1.05*10^{-4}$-$5.22*10^{-4}$μg/mL (21) | ~10μg/ml |

**Supplementary Table 9.** Summary of growth factor sources and concentrations used in this study, with reference to physiological ranges and manufacturer specifications.

**Supplementary Table 10.**

| Compound | 3.3µM | 1.1µM | 0.37µM | 0.12µM | 0.04µM | 0.01µM |
|---|---|---|---|---|---|---|
| Cdk1/2 inhibitor III | xx | xx | xx | xx | ✓✓ | - |
| LCK inhibitor RK24466 | - | - | ✓✓ | ✓✓ | - | ✓✓ |
| PKC412 | xx | xx | xx | - | ✓✓ | - |
| JAK inhibitor VI | xx | xx | - | - | - | - |
| Vargatef | xx | xx | xx | xx | xx | xx |
| NCGC00244250-01 | xx | xx | - | - | xx | - |
| AMG-51 | xx | xx | xx | - | - | - |
| Go 6976 | xx | xx | xx | - | - | - |
| PRT-060318 | xx | xx | xx | - | - | - |
| Bosutinib | xx | xx | - | xx | xx | xx |
| NVP-BGT226 | xx | xx | xx | xx | xx | xx |
| JNJ-7706621 | xx | - | ✓✓ | ✓✓ | - | - |
| WZ3416 | xx | xx | - | - | - | - |
| GSK650394 | - | - | - | - | - | - |
| AT7867 | - | ✓✓ | ✓✓ | ✓✓ | ✓ | - |
| Motesanib | - | - | xx | xx | xx | - |
| PD-173955 Analogue 1 | - | ✓✓ | - | - | - | - |
| PP121 | xx | xx | xx | xx | xx | |
| Pluripotin | xx | xx | xx | xx | xx | xx |
| PF-3758309 | xx | xx | xx | xx | - | - |
| Neratinib | xx | xx | xx | xx | - | - |
| Tivozanib | xx | xx | xx | - | - | - |
| Dasatinib | - | - | - | - | - | - |
| Crenolanib | xx | - | - | xx | - | xx |
| TWS119 | xx | xx | xx | - | - | - |
| AT9283 | xx | xx | xx | xx | xx | xx |
| BMS-754807 | xx | xx | xx | xx | xx | xx |
| Erlotinib hydrochloride | - | - | - | - | - | - |
| PHA-848125 | xx | xx | xx | xx | xx | xx |
| Dinaciclib | xx | xx | xx | xx | xx | xx |
| DCC-2036 | xx | xx | xx | xx | xx | xx |
| Roscovitine | xx | xx | - | - | xx | - |
| Bisindolylmaleimide | - | xx | xx | xx | xx | xx |
| NCGC00348110 | - | - | - | xx | xx | - |
| BIIB-057 | - | xx | - | - | - | - |
| AP26113 | xx | xx | x | - | - | - |

| | | | | | | |
|---|---|---|---|---|---|---|
| MGCD-265 | xx | xx | xx | xx | xx | xx |
| GSK-269962A | ✓✓ | ✓✓ | ✓✓ | ✓✓ | - | - |
| GZD824* | xx | xx | xx | xx | xx | xx |
| Ponatinib | xx | xx | xx | xx | xx | - |
| LDN 193189 | xx | xx | - | - | - | - |
| CHEMBL2336015 | xx | - | - | - | - | - |
| TAE226 | xx | xx | xx | xx | xx | - |
| PF-3644022 | - | x | - | - | - | xx |
| PKI-SU11274 | - | - | ✓✓ | ✓✓ | ✓✓ | ✓✓ |
| Regorafenib | xx | xx | xx | - | x | xx |
| Lestaurtinib | xx | xx | xx | xx | xx | xx |

**Supplementary Table 10.** Treatment effects on anastasis across all post-wash time points. (xx) indicates -1, (x) -0.5, (-) 0, (✓) +0.5, (✓✓) +1. *3.33 and 1.11 µM GZD824 have been corrected by manual counting (see Methods Section 4.9.1).
